# Low-bias ncRNA Libraries using Ordered Two-Template Relay: Serial Template Jumping by a Modified Retroelement Reverse Transcriptase

**DOI:** 10.1101/2021.04.30.442027

**Authors:** Heather E. Upton, Lucas Ferguson, Morayma M. Temoche-Diaz, Xiaoman Liu, Sydney C. Pimentel, Nicholas T. Ingolia, Randy Schekman, Kathleen Collins

**Affiliations:** Department of Molecular and Cell Biology, University of California at Berkeley, Berkeley, CA 94720; Bakar Innovation Fellow, University of California at Berkeley; Computational Biology Program, University of California at Berkeley; Department of Plant and Microbial Biology, University of California at Berkeley; Howard Hughes Medical Institute; Bakar Fellow, University of California at Berkeley

**Keywords:** non-LTR retroelement reverse transcriptase, RNA sequencing, miRNA, tRNA, non-coding RNA

## Abstract

Non-long terminal repeat (non-LTR) and group II intron retroelements encode reverse transcriptases (RTs) that copy the retroelement transcript directly into host cell DNA, often at specific target sites. Biochemical characterization of these enzymes has been limited by recombinant expression and purification challenges, hampering understanding of their transposition mechanism and their exploitation for research and biotechnology. Properties of retroelement RTs substantiate their application for end-to-end RNA sequence capture. To investigate this utility, we first compared a non-LTR RT from *Bombyx mori* and a group II intron RT from *Eubacterium rectale*. Only the non-LTR RT showed processive template jumping, producing one cDNA from discontinuous templates each copied end-to-end. We also discovered an unexpected terminal deoxynucleotidyl transferase activity of the RTs that adds nucleotide(s) of choice to 3’ ends of single-stranded RNA or DNA. Combining these two types of activity with additional insights about non-templated nucleotide additions to duplexed cDNA product, we developed a streamlined protocol for linking Next Generation Sequencing (NGS) adaptors to both cDNA ends in a single RT reaction. When benchmarked using a reference pool of microRNAs (miRNAs), library production using modified non-LTR retroelment RT for Ordered Two-Template Relay (OTTR) outperformed all commercially available kits and rivaled the low bias of technically demanding home-brew protocols. We applied OTTR to inventory RNAs purified from extracellular vesicles (EVs), identifying miRNAs as well as myriad other non-coding (nc) RNAs and ncRNA fragments. Our results establish the utility of OTTR for automation-friendly, low-bias, end-to-end RNA sequence inventories of complex ncRNA samples.

**Significance:** Retrotransposons are non-infectious mobile genetic elements that proliferate in host genomes via an RNA intermediate that is copied into DNA by a reverse transcriptase (RT) enzyme. RTs are important for biotechnological applications involving information capture from RNA, since RNA is first converted into complementary DNA for detection or sequencing. Here, we biochemically characterize RTs from two retroelements and uncover several activities that allowed us to design a streamlined, efficient workflow for determining the inventory of RNA sequences in processed RNA pools. The unique properties of non-retroviral RT activities obviate many technical issues associated with current methods of RNA sequence analysis, with wide applications in research, biotechnology, and diagnostics.

## Introduction

Retroelements are mobile genome segments that use an RNA intermediate to template the synthesis of DNA inserted at a new genome location. This group of selfishly replicating DNAs includes eukaryotic long terminal repeat (LTR) and non-LTR retrotransposons, as well as prokaryotic mobile introns also found in eukaryotic organelles. Genome sequencing projects have revealed evolutionary episodes of dramatic retroelement proliferation, for example the spread of human non-LTR long interspersed nuclear element 1 (LINE-1) to constitute about 20% of our genome (1). Many non-LTR retroelements in the genomes of living organisms retain the ancestral eukaryotic retroelement architecture (2), with a single open reading frame (ORF) between unique 5′ and 3′ untranslated regions (UTRs). Not surprisingly, the element-encoded protein multitasks in its interactions with RNA and DNA, reverse transcriptase (RT) activity, and often DNA-nickase activity. Some of these retroelements show site-specific insertion, which would limit their copy number, decrease their toxicity, and increase their potential for long-term evolutionary persistence (3).

The only detailed biochemical characterization of a purified RT from the ancestral single-ORF non-LTR retroelement families is of the R2 element protein from the silkmoth *Bombyx mori*, in pioneering work by the Eickbush laboratory (4). R2 and other R-elements insert exclusively into a precise sequence of the ribosomal RNA (rRNA) precursor gene (rDNA) transcribed by RNA Polymerase I (RNAP I) (5). The R2 ORF encodes a protein comprised of N-terminal DNA-binding motifs (zinc-finger and Myb domains), a central RT domain, a C-terminal restriction-like endonuclease domain (EN), and other regions of unknown function (Fig. S1A). After binding the target site and introducing a nick to create a DNA primer 3′ end, the protein then switches to reverse transcription of a bound template RNA to produce complementary DNA (cDNA). This process is termed target-primed reverse transcription (TPRT) (4). To complete non-LTR retroelement insertion, second-strand DNA nicking and synthesis must also occur, the latter of which could in theory be performed by the retroelement protein and/or a cellular DNA polymerase (4).

Recombinant *B. mori* R2 protein produced in bacteria, combined with target DNA duplex and an RNA containing the retroelement 3′ UTR, is sufficient to reconstitute site-specific TPRT in vitro (6). The RT initiates cDNA synthesis by “template jumping,” which we define as engaging the 3′ end of an RNA to template cDNA synthesis without base-pairing or with just 1-2 base-pairs that would only be stable within the enzyme active site. Group II intron RTs are known to template-jump in vitro, but in cells, intron insertion begins by reverse-splicing of the catalytic RNA followed by cDNA synthesis on the contiguous DNA-RNA template (7). Retroviral RTs have template-jumping activity in vitro exploited for cDNA library 3′ adaptor addition (8), but the required amount of base pairing between cDNA and template RNA is uncertain (9). Template jumping differs from retroviral RT “template switching,” which we define as occurring by cDNA product release from one template and re-annealing to a new template anywhere within a transcript. This template switching activity is essential to complete synthesis of LTRs (10).

In principle, the use of template jumping to make cDNA libraries that capture template sequences end-to-end, flanked by 5′ and 3′ adaptors, could be a boon for research and biotechnological applications. New methods of RNA sequencing (RNA-seq) have illuminated an ever-increasing diversity of RNA types, but challenges associated with library generation from non-coding RNA (ncRNA) limit what is known about specifically processed forms of ncRNA in cells and in extracellular vesicles (EVs) (11–14). Towards the goal of comprehensive, unbiased, end-to-end ncRNA-seq, we sought to use retroelement RT(s) for serial template jumping to add distinct 5′ and 3′ adaptors during cDNA library synthesis. We first compared RTs from group II introns and non-LTR retroelements for maximal template-jumping activity, and then we engineered the rampant template-jumping of a truncated, modified R2 RT to perform two template jumps in specific order. We describe streamlined, automation-friendly, single-tube cDNA library production for next-generation sequencing (NGS) with library indexing by the initial cDNA library synthesis or by low-cycle PCR. We benchmarked the new technology by sequencing cDNA libraries produced from a commercial reference standard of 962 microRNAs (miRNAs). Next, to gain biological insight, we used the technology to sequence small RNAs (sRNAs) in EVs secreted by human cell lines, with results that have implications for models of EV biogenesis and function.

## Results

### Comparison of template jumping by non-retroviral RTs

We evaluated the ability of non-retroviral RTs to use physically separate, discontinuous template molecules for continuous cDNA synthesis (primer and template sequences are listed in Table S1). We screened recombinant versions of bacterial intron and eukaryotic non-LTR RTs for robust expression, purification, and serial template jumping. Proteins were expressed as N-terminal maltose binding protein (MBP) fusions with a C-terminal 6-histidine (6xHis) tag. Among the group II intron RTs tested under pilot screen conditions, the top candidate was from *Eubacterium rectale* (EuRe, Figs. 1 and S1A). In papers published subsequent to our initial screening, this same enzyme, differing in tag configuration, was shown in studies from the Pyle laboratory to support remarkably processive cDNA synthesis (15, 16). Group II intron RTs synthesize cDNA across base- and sugar-modified templates with high tolerance for RNA structure (17), contributing to their processivity. In our assays of serial template jumping, the best performing enzyme was an N-terminally truncated eukaryotic R2 non-LTR retroelement RT from *B. mori* (BoMoC, Figs. 1 and S1A). Because removal of the C-terminal EN domain of BoMoC was unfavorable for enzyme stability, we introduced an EN active-site mutation previously characterized in the full-length protein (18) to produce an endonuclease-dead version of BoMoC. This modified, truncated R2 RT enzyme was used for all experiments described below.

**Figure 1.**
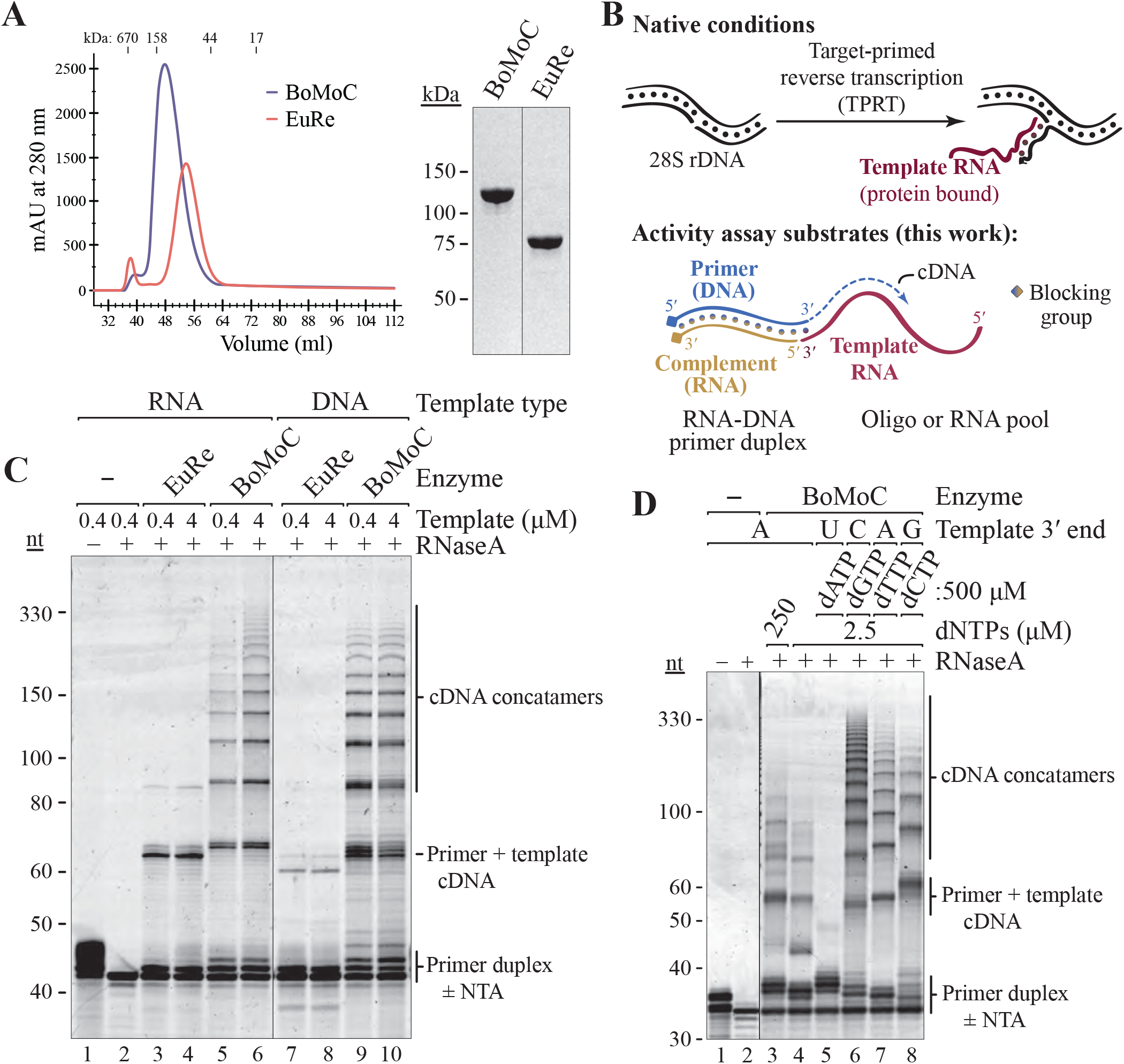
Recombinant cellular retroelement RTs differ in template jumping processivity. A) Size-exclusion chromatography (left) and SDS-PAGE (right) of purified BoMoC and EuRe RTs. B) Schematic of non-LTR retrotransposon cDNA synthesis by TPRT (top) and cDNA library synthesis by OTTR (bottom). C) SYBR Gold-stained denaturing PAGE gel of BoMoC and EuRe RT products using +1T primer duplex and a DNA or RNA oligonucleotide template. D) SYBR Gold-stained denaturing PAGE gel of BoMoC RT products using blunt-end primer duplex with dNTPs and RNA templates indicated.

Recombinant EuRe and BoMoC RTs were purified extensively to remove nucleic acid and nuclease contamination. Purification involved binding to and elution from nickel agarose and heparin agarose, with a final step of size-exclusion chromatography (SEC), a strategy also used in previous EuRe purifications (16, 19). Very high ionic strength buffers were essential to release bound nucleic acids and decrease aggregation. Both EuRe and BoMoC fractionated as monomers by SEC and migrated in SDS-PAGE consistent with predicted fusion proteins of 91 kDa and 137 kDa, respectively (Fig. 1A). Measurement of the absorbance ratio at 260 and 280 nm suggested a lack of nucleic acid co-purifying with the RT proteins, as did attempts to detect bound nucleic acid by SYBR Gold staining (data not shown). MBP tag removal from bacterially produced BoMoC reduced its stability; therefore, the RTs were used as fusion proteins for all assays. We have detected no obvious activity difference between bacterially expressed MBP-BoMoC-6xHis and BoMoC expressed without an MBP tag purified from human cells (data not shown).

We assayed template jumping using an oligonucleotide primer duplex and single-stranded template, mimicking the physiological process of TPRT (Fig. 1B, top). We designed a primer duplex with a 3′ OH DNA strand and a 3′-blocked RNA strand such that only the DNA strand could be elongated (Fig. 1B, bottom). BoMoC synthesized cDNA products by continuous primer extension across at least 10 template molecules. This highly processive template jumping was observed in reactions with RNA or DNA templates (Fig. 1C). Curiously, in comparison to BoMoC, EuRe had relatively weak template jumping activity and strong preference for RNA as template (Fig. 1C).

To investigate the influence of a DNA primer 3′ overhang on template jumping, we used primer duplexes that differed only in length of the DNA 3′ overhang. More than 2 nucleotides (nt) of overhang was strongly inhibitory for cDNA synthesis by BoMoC, even if templates had a 3′ sequence fully complementary to the primer 3′ overhang (Fig. S1B). We next tested how overhang sequence affected template choice by comparing use of a +1C or +1T overhang primer for jumping to the same set of templates. The +1T primer supported cDNA synthesis from a template with 3′A but not 3′G, whereas the +1C primer supported cDNA synthesis from 3′G and less efficiently also 3′A templates (Fig. S1C). However, products from the +1C primer using a template with 3′A were slightly shorter than products from reactions using the +1T primer and the same template, indicative of the +1C overhang pairing with the template G 1 nt internal to the template 3′ end. The ability of a G-C base-pair to allow for initiation slightly internal to the template 3′ end was previously noted for full-length BoMo R2 RT protein under conditions when the enzyme can not make an appropriate base-pairing of primer overhang and template 3’ end (20). From these results we conclude that use of a primer containing a +1 overhang at least partially suppresses use of templates with a non-complementary 3′ end. Also, primer 3’ overhangs of >2 nt are inhibitory for template jumping. BoMoC preferred a 1 nt overhang versus 2 nt overhang, opposite the preference of a retroviral RT (21).

### Relationship between non-templated nucleotide addition and template jumping

DNA-templated DNA polymerases, especially those without a 3′-5′ proofreading exonuclease activity, tend to dissociate after adding a single-nt overhang to a cDNA duplex (22–25). BoMoC, as an RNA- or DNA-templated polymerase, adds several non-templated nt to a fully duplexed primer or product 3’ end. This can be clearly visualized in the primer-extension products 1-5 nt longer than the starting primer (Figs. 1C, 1D, and S1D). The addition of a typically 3-4 nt 3′- overhang, with some product having a 5 nt 3’-overhang, suggests that BoMoC may have even more robust non-templated nt addition (NTA) than the full-length R2 RT protein shown to add a 2-3 nt 3′ overhang (20). This difference could be inherent to the protein sequences or, more likely, differences in enzyme purification, storage, and reaction conditions.

NTA could facilitate template jumping by creating a cDNA 3′ overhang that base-pairs to a template 3′ end. On the other hand, NTA could be the consequence of aborted template jumping rather than a stimulus for it. For retroviral RTs, different studies come to different conclusions about NTA dNTP preference and number of nt added, as well as the role of NTA in template jumping (9, 21, 26–29). For BoMoC, we first compared NTA and template jumping activities in the presence of a 200-fold excess of each single dNTP over the other dNTPs using a blunt-end primer duplex and templates with a 3′ nt complementary to the dNTP in excess (Fig. 1D). Reactions with a high dATP concentration promoted maximal NTA, as noted with full-length R2 RT (20). Template jumping in this high dATP reaction was dramatically suppressed, yielding almost only products corresponding to primer extended by +3 and +4 NTA (Fig. 1D, lane 5). In comparison, reactions with excess dGTP allowed 2-3 nt of NTA and maximal template jumping (Fig. 1D, lane 6). Reactions with excess dCTP or dTTP generated products with typically 2 nt of NTA and intermediate template jumping processivity. Together, these assays do not point to a simple relationship between efficiency of NTA and template jumping. We suggest that the +2 to +5 NTA products that accumulate are inhibitory to template jumping, whereas the low level of +1 NTA product in part reflects its use for additional cDNA synthesis by template jumping. From this perspective, NTA is stimulatory for template jumping but only under conditions that limit or slow extension of a +1 nt overhang to +2 nt.

Curiously, in reactions using primers with a +1T 3’ overhang, we observed very little NTA to extend the +1 overhang even if the same dNTP concentration induced up to 5 nt of NTA on a blunt-end primer duplex (Fig. S1D, compare the NTA products of the primer). The reduction of NTA using a +1T primer appeared to promote the initial template jump, particularly in reactions with equal concentrations of each dNTP (Fig. S1D; note that only the first template jump would be influenced by a primer 3’ overhang). Qualitatively similar results were observed using primers with +1C and +1G but not +1A (data not shown). This led us to develop a strategy for ordered serial template jumping dependent on a +1T RNA-DNA duplex to capture the first template (see below).

### Terminal transferase activity in the presence of manganese ions

Polymerases require divalent cations for catalysis. Typically, Mg^2+^ functions as the cofactor under physiological conditions but other divalent ions, including Mn^2+^, can support some level of DNA synthesis. To determine how BoMoC activity is influenced by use of Mn^2+^, we substituted Mn^2+^ for Mg^2+^ in template-jump reactions. Expected cDNA products were not detected; instead, a smear of variable length product was observed. Surprisingly, in reactions with Mn^2+^, BoMoC added non-templated nt(s) to the 3′ end of single-stranded RNAs or DNAs and also to double-stranded RNA-RNA, DNA-DNA, or RNA-DNA substrates (Fig. 2A). This type of activity is often described as terminal transferase or “tailing” activity (30, 31). Assays of EuRe for Mn^2+^- dependent terminal transferase activity showed it to have less tailing activity than BoMoC (Fig. S2A). Previous studies have shown that full-length BoMo R2 RT can use single-stranded RNA to prime synthesis across another non-complementary oligonucleotide or transcript (32). We observed some products in Mg^2+^ reactions with BoMoC and EuRe that likely arise from this non-selective priming (Figs. 2A and S2A, asterisks). Unfortunately, cross-priming of single-stranded RNA or DNA molecules intended to be template molecules compromises the template pool by depletion and synthesis of artifact chimeric products.

**Figure 2.**
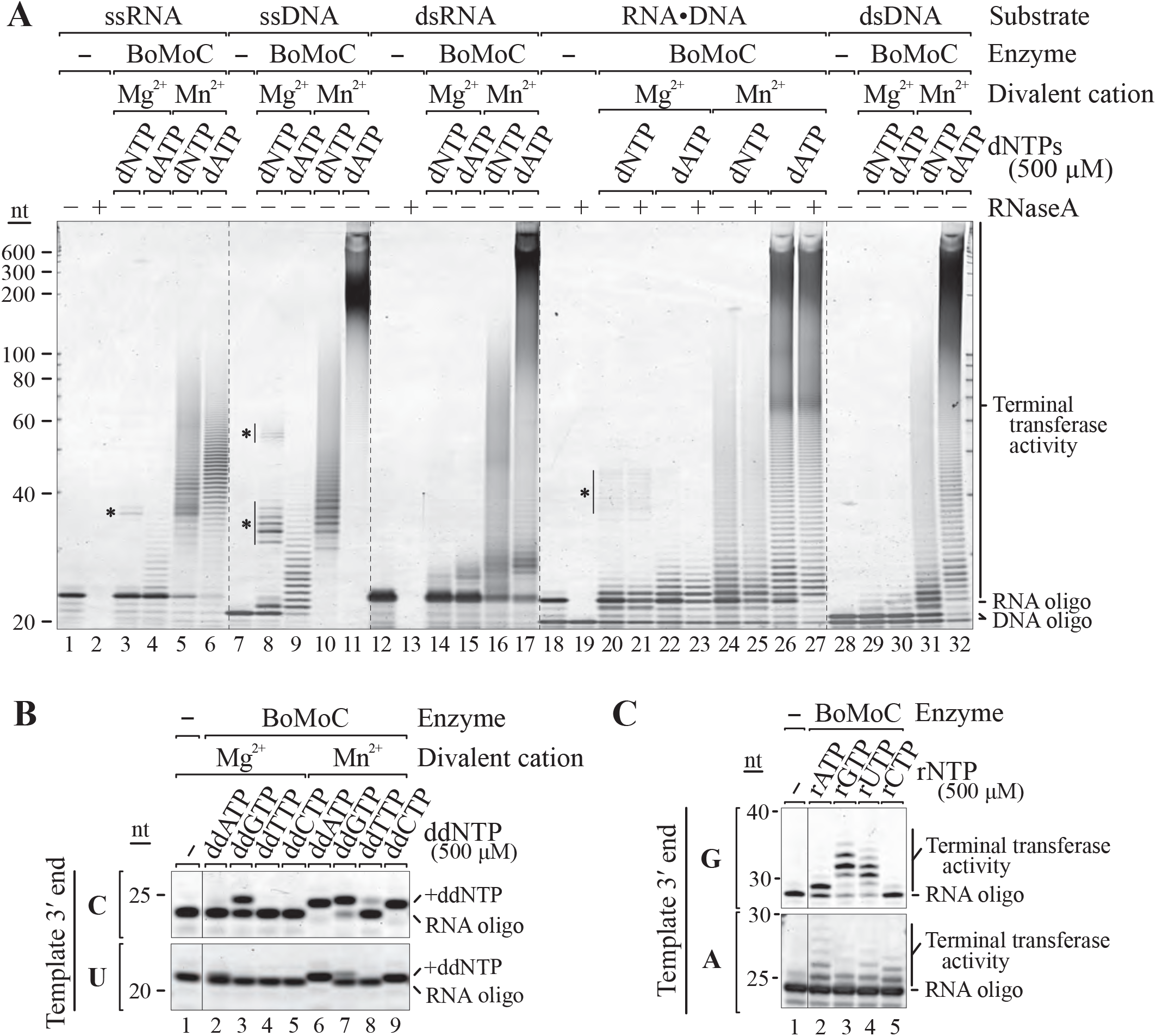
BoMoC acts as a terminal transferase in the presence of Mn^2+^. SYBR Gold-stained denaturing PAGE gels of the terminal transferase reaction products of BoMoC are shown. A) Input single-stranded (ss) RNA and DNA and double-stranded (ds) blunt-ended RNA, DNA, and RNA-DNA duplex were assayed in Mg^2+^ and Mn^2+^ reaction conditions in the presence of 500 μM of each dNTP or dATP alone. Products marked with asterisks indicate template copying primed by a non-complementary oligonucleotide. B) RNA oligonucleotide with 3’C or 3’U was assayed for extension by a single ddNTP in Mg^2+^ and Mn^2+^ conditions, in the presence of 500 μM of an individual ddNTP. Single nucleotide addition in Mg^2+^ results from cDNA synthesis priming by a non-complementary oligonucleotide. C) RNA oligonucleotide with 3’G or 3’A was assayed for extension under Mn^2+^ conditions in the presence of 500 μM of an individual rNTP.

In Mn^2+^ reactions with a single dNTP, BoMoC added a dNTP-dependent length of homopolymer tract (Figs. S2B and S2C). Tailing of single-stranded RNA or DNA by incorporation of dATP was especially rampant compared to tailing with mixed dNTPs or other individual dNTPs (Figs. 2A, S2B, and S2C). Studies of the Tf1 *Schizosaccharomyces pombe* LTR-retroelement RT demonstrated tailing in Mn^2+^ specifically with dATP (33), although this activity seems much less processive than tailing by BoMoC. In single-stranded RNA tailing reactions, BoMoC had substrate preference related to the primer 3′ nt. For example, a substrate ending with 3′G generally showed compromised tailing with dCTP, whereas a template with 3′A generally showed compromised tailing with dTTP (Figs. S2B and S2C). We suggest that in Mn^2+^ reaction conditions, a single-stranded nucleic acid will preferentially bind as template in the active site in the presence of a 3’-end cognate dNTP, whereas with a non-cognate dNTP it will more readily bind as primer to enable its extension by terminal transferase activity.

To investigate whether BoMoC could give a pool of template nucleic acids a single, shared 3′ nt, we assayed incorporation of dideoxynucleotides (ddNTPs) in tailing reactions. Each ddNTP could be incorporated to some extent (Fig. 2B). As observed in reactions with dNTPs, addition of ddNTPs was similarly influenced by the template 3′ nt. The most efficient and general labeling was observed in reactions with ddATP and was improved for difficult substrates by lower reaction temperatures (30°C compared to 37°C), the presence of a crowding agent (PEG-8000), and increased reaction time (Fig. S2D). A limited extent of tailing by ribonucleotide addition was also observed (Fig. 2C), even without an active-site mutation to remove steric hindrance on the ribose 2′ hydroxyl (34).

### Ordered two-template relay for cDNA library synthesis

NGS libraries from sRNA are commonly prepared by sequential ligation of adaptors to input RNA 3′ and 5′ ends, often with a gel purification step after each ligation step, followed by cDNA synthesis and PCR. Low bias in miRNA capture is possible using days-long home-brew protocols with degenerate sequence adaptor ends (35). However, this type of protocol is time consuming in manual effort, technically challenging, and requires high input RNA due to many steps with product loss. We sought to exploit the serial template jumping ability of BoMoC as the basis of a ligation-independent method for end-to-end sRNA library synthesis for NGS.

Ultimately, we developed Ordered Two-Template Relay (OTTR): a single-tube reverse transcription reaction for dual-end adaptor-tagged cDNA library synthesis (Fig. 3A). First, we used the terminal transferase activity of BoMoC to add a single ddRTP (ddATP and/or ddGTP) to input template (IT) RNA 3′ ends (Fig. 3A, maroon line). By utilizing primer duplex(es) with a +1Y (+1T and/or +1C) overhang (Fig. 3A, blue/tan duplex), all IT molecules could form a single base-pair between template and primer 3′ ends. This strategy exploits our observation that a primer +1Y overhang is particularly resistant to additional NTA that would inactivate the primer for template jumping. In the same RT reaction, we added a template for synthesis of a single copy of cDNA 3′ adaptor. To disfavor use of this cDNA 3’ adaptor template (AT) until after cDNA synthesis across an IT, the AT has a 3′C (Fig. 3A, green line). By manipulating dNTP concentrations and adding a dNTP analog to the reaction, we encouraged extension of the IT cDNA by a single NTA of dGTP. This gives the AT, but not IT, an ability to form a single base-pair with the intermediate-stage cDNA 3’ overhang, recruiting the 3′C AT for the second template jump to complete library synthesis. If the AT has a 5′ block to additional template jumping, the desired cDNA library is produced. Adaptor dimer formation is limited by the mismatch between primer +1Y and AT 3′C. Copying of more than one molecule of input sRNA is strongly suppressed by the extremely poor use of a dYTP for NTA to the intermediate-stage cDNA and by poor elongation of a mismatched cDNA 3’G by template jumping to an IT with a 3’ ddR. Importantly, 3’ tailing of input sRNA with a non-extendable ddNTP prevents artifact generation by hybridization-independent template priming of cDNA synthesis on another input RNA (36, 37), which would deplete the template pool and generate non-native fusions.

**Figure 3.**
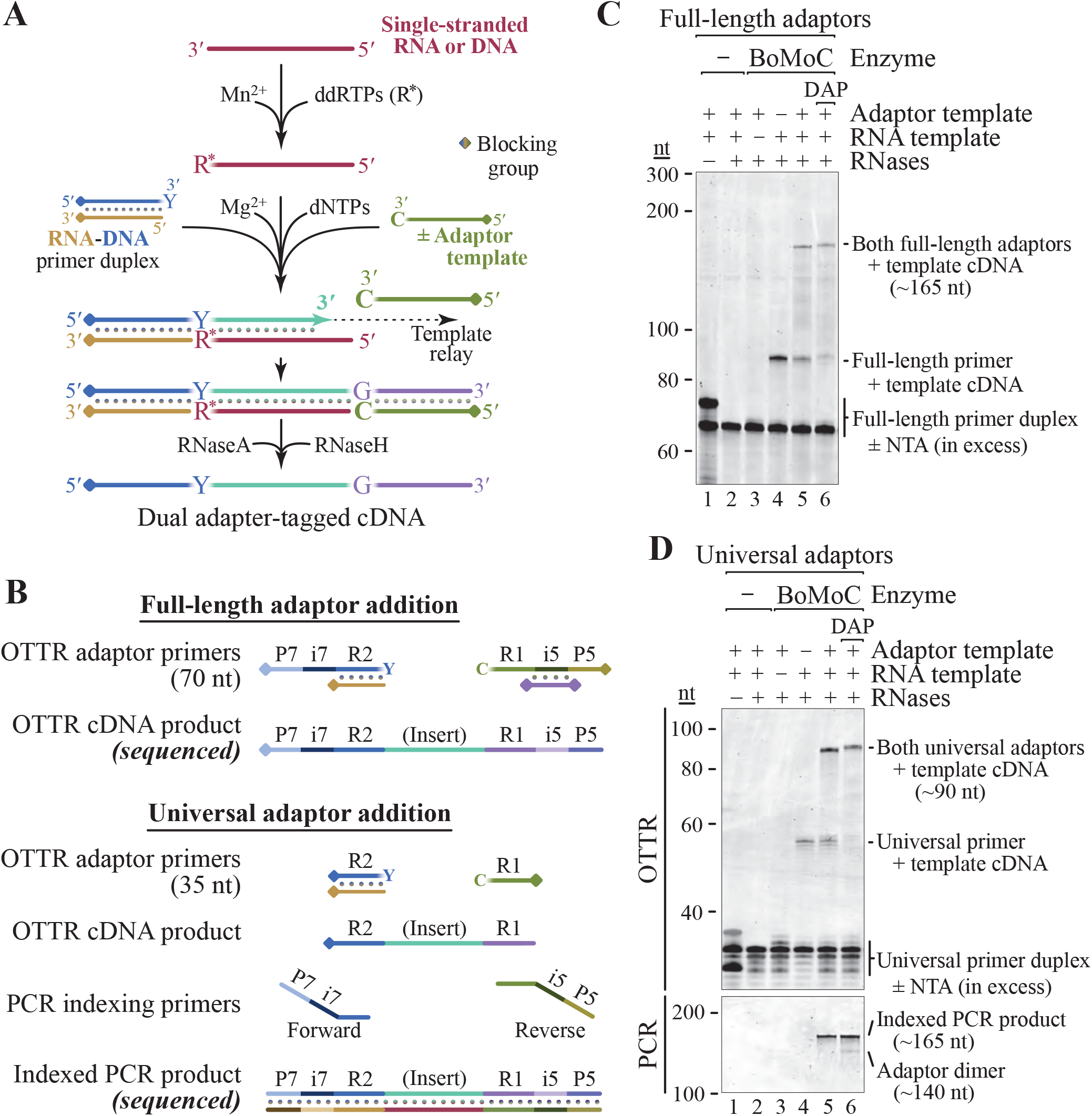
OTTR for NGS cDNA library generation. A) Optimized workflow for single-tube synthesis of cDNA libraries. A pool of RNA and/or DNA input molecules (maroon) is first labeled with 3’ ddRTP. Subsequently free ddRTP is inactivated and buffer conditions are toggled from Mn^2+^ to Mg^2+^. Next, dNTPs, adaptor oligonucleotides, and BoMoC are added to initiate cDNA synthesis from the RNA-DNA primer duplex across the IT (maroon), ending after copying the AT (green). Products are then treated with RNaseA and RNaseH to remove RNA, yielding the desired cDNA. B) Schematic of primers involved in Illumina Full-length (top) or Universal (bottom) adaptor addition and their respective cDNA library products. DNA primers were the complement of P7-i7-R2 or R2, while ATs were P5-i5-R1 or R1. In the full-length adaptor strategy, only cDNA products elongated by copying the AT can bind to the flow cell. C, D) Proof of principle for OTTR library generation using an RNA oligonucleotide template with Full-length (C) or Universal (D) adaptors. Only reactions containing primer duplex, RNA template, adaptor template, and BoMoC (lanes 5 and 6) generate properly sized cDNA library product. Universal-adaptor RT reactions required PCR amplification for P5 or P7 sequence fusion and indexing (D, bottom). DAP: diaminopurine deoxyribose triphosphate.

In the OTTR protocol, 5′ and 3′ cDNA adaptor sequences can be varied as desired. We confirmed cDNA library synthesis using both the Illumina NGS “Universal” read 1 (R1) and read 2 (R2) adaptor sequences and the “Full-length” P5-i5-R1 and P7-i7-R2 adaptor sequences (Fig. 3, Table S1). The Universal adaptor cDNA libraries were indexed by low-cycle PCR (4-8 cycles) with P5-i5 and P7-i7 primers, whereas the Full-length adaptor cDNA libraries were indexed by inclusion of different i5 and i7 bar codes in the 5′ cDNA primer and 3′ cDNA AT included in the RT reaction (Fig. 3B). In some experiments, we placed a unique molecular identifier in the 3′ AT oligonucleotide, adding 5 or more random nucleotides (N) adjacent to a YC-3′ end with no change in library yield or bias (data not shown).

We optimized OTTR using RNA oligonucleotide templates. The intended dual-adaptor-flanked cDNAs were generated when all reaction components were present (Figs. 3C and 3D). In our initial workflows, the yield of complete cDNA library product was diminished by some +2 or more nt of NTA that inactivates the intermediate cDNA for the second template jump (Figs. 3C and 3D, lane 5). We found that replacement of most of the dATP in the reaction with diaminopurine deoxyribose triphosphate (DAP) nearly eliminated the dead-end intermediate cDNA products (Figs. 3C and 3D, lane 6), presumably due to reduced tailing.

### OTTR outperforms commercial kits for making NGS miRNA libraries

To evaluate OTTR for ncRNA library preparation for NGS, we tested OTTR with the miRXplore reference standard. This reference contains a reported 963 distinct synthetic miRNAs at equimolar ratio (in fact 962 as two sequences are identical). Each RNA oligonucleotide has a 5′ monophosphate and 3′ hydroxyl group like a native miRNA. Many commercially available cDNA library kits have been evaluated using the miRXplore reference standard, enabling us to sample independently obtained data to benchmark OTTR against commercial kits (35, 38–40).

Quantitative evaluation of cDNA library capture bias of miRNAs in the miRXplore standard for each method can be done by calculating the coefficient of variation (CV), defined as the ratio of standard deviation to mean read counts totaled across all input sequences in a sample. If individual miRNAs have read counts far from the expected mean, the CV increases; therefore, the lower the CV, the better the library. The same read count information can be visualized as a violin plot of miRNA read counts, with each miRNA adding to violin width or height on the vertical axis of read counts (Fig. 4A). A good library has a short and wide violin, indicating that most miRNAs had read counts close to expected. In addition to CV, read-count violin plots, and the number of miRNAs detected per fixed number of reads, we evaluated bias by clustering libraries according to similarities of bias for each individual miRNA (Fig. 4A).

**Figure 4.**
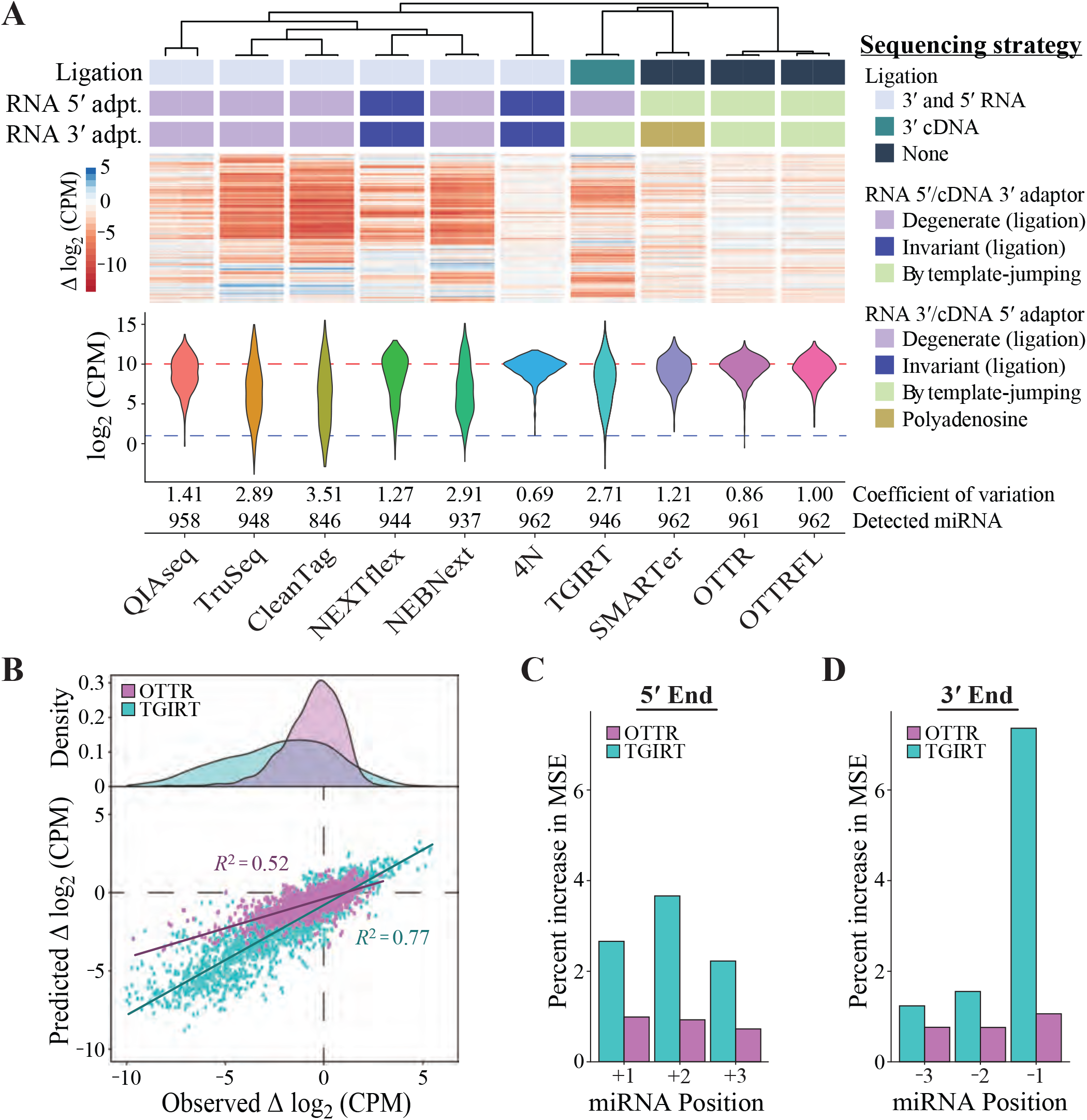
OTTR outperforms library generation protocols from commercially available kits. A) Unsupervised hierarchical clustering of Δlog_2_CPM of miRNA read counts from libraries made using different protocols, with side-by-side paired technical replicates for each. Δlog_2_ CPM = log_2_ (CPM_expected_ / CPM_observed_), where CPM_expected_ is 962^−1^ × 1,000,000. The dendrogram indicates the relatedness of miRNA read-count bias. Annotations below the dendrogram indicate protocol distinctions in ligase and polymerase usage. Ligated adaptors were considered “degenerate” or “invariant” based on whether the adapter sequence had mixed-base positions. “Polyadenosine” indicates tailing of the input RNA by polyA polymerase necessary for binding of an oligothymidine RT primer. DESeq2 was used to normalize read counts for each set of replicates before conversion to log_2_ CPM, and the distributions for combined replicates are presented as violin plots. Across the violins, the red dashed line defines the expected mean log_2_CPM of equimolar representation and the blue dashed line defines the detected cutoff, which was CPM > 2. B). Evaluation of random forest models’ predicted Δlog_2_CPM and observed Δlog_2_ CPM for each miRNA based on the 5’ most (+1, +2, and +3) or 3’ most (−3, −2, −1) bases. C, D). Percent increase in mean squared error (MSE) or relative importance for each variable of the random forest model trained on OTTR and TGIRT datasets (C: +1, +2, and +3 for 5’ three-most bases where +1 is the exact 5’ end; D: −3, −2, and −1 for 3’ most bases where −1 is the exact 3’ end). Variables with a higher percent MSE are considered more important in the random forest model when predicting the log_2_ CPM.

We randomly sampled a matched number of deposited reads from Illumina NGS libraries of the miRXplore standard and reads from two types of OTTR libraries: one with Universal cDNA adaptors and indexing by PCR (“OTTR”) and one with Full-length NGS cDNA adaptors and no PCR (“OTTRFL”). The CVs of OTTR libraries were lower than all commercial kits, and the violin plots showed more miRNAs with read-counts near the expected log_2_-scale count per million of 10 (Fig. 4A, bottom). The lowest CV among the sampled cDNA libraries was observed for an extensively optimized, ligation-based, home-brew protocol with a technically challenging workflow (“4N”). The ligation-based kits clustered in their profiles of miRNA read count deviation from equal representation, as did OTTR protocols with or without PCR (Fig. 4A, top). While we used this benchmarked version of OTTR for ncRNA discovery studies described below, ongoing improvements made the OTTR miRXplore miRNA library CV lower than that of 4N-protocol libraries (data not shown).

We were particularly interested in comparison of miRNA capture bias between OTTR and other protocols that include some kind of template jumping. TGIRT-seq (41, 42) uses a bacterial thermostable intron RT, TGIRT, for an intended single template jump to initiate cDNA synthesis (i.e. to jump “on” an input template RNA). SMARTer-seq (43) uses a modified retroviral RT for an inefficient single template jump to extend the initial cDNA by synthesis across an adaptor template (i.e. to jump “off” the duplex of input template and its cDNA). Only OTTR exploits serial ordered template jumping to add both adaptors in a single step. The bias of TGIRT-based library generation is substantially higher than that of OTTR (Fig. 4A). Comparison of TGIRT-seq to OTTR using a scatter plot of expected versus observed individual miRNA read counts offers a granular visualization of their difference (Fig. 4B). TGIRT-seq bias appears to derive predominantly from the identity of the template 3′ nt that would engage the +1N primer overhang to support template jumping, with less bias from the template 5′ end (Figs. 4C and 4D). Although the SMARTer protocol has the best CV among commercial kits, it is also outperformed by OTTR (Fig. 4A) and is compromised in utility by the loss of sRNA 3’ end information due to polyadenosine tailing prior to dT-primed cDNA synthesis.

### OTTR for EV RNA sequencing

Many categories of sRNA, including miRNA, piwi-interacting RNA (piRNA), tRNA/tRNA fragments (tRFs), and Y RNAs, play important roles in the regulation of gene expression (44, 45). The profile of these RNAs in the bloodstream and other biofluids holds promise as an approach for diagnostic monitoring of human disease (46, 47). Extracellular RNAs with more than a fleeting half-life are contained within EVs, a vesicle population that includes low-density EVs released by budding from the plasma membrane and higher-density EVs released upon fusion of cytoplasmic multivesicular bodies with the plasma membrane (48).

To improve community knowledge of EV sRNA inventories, we generated and sequenced OTTR cDNA libraries from EV populations. From the breast-cancer derived MDA-MB-231 cell line, EVs were sampled as crude EV preparations from conditioned medium by single-step centrifugation (100,000 x g pellet containing vesicular and non-vesicular sedimentable material) and as highly purified vesicles floated in a sucrose density step gradient (Floated EVs) and treated with micrococcal nuclease (MNase) before detergent lysis to eliminate any nucleic acids not enclosed within the vesicles (Fig. 5A). The length profile of total cellular RNA included major peaks for 18S and 28S rRNAs and tRNAs, whereas bulk or highly purified EV RNAs had lengths predominantly of tRNA size or smaller (Fig. 5B). For sequencing comparison we used filtration to enrich total cellular sRNA of less than 200 nt prior to library generation. We also used a different human cell line, HEK 293T, to generate similarly size-enriched total cellular sRNA and floated vesicles for comparison (Fig. S3).

**Figure 5.**
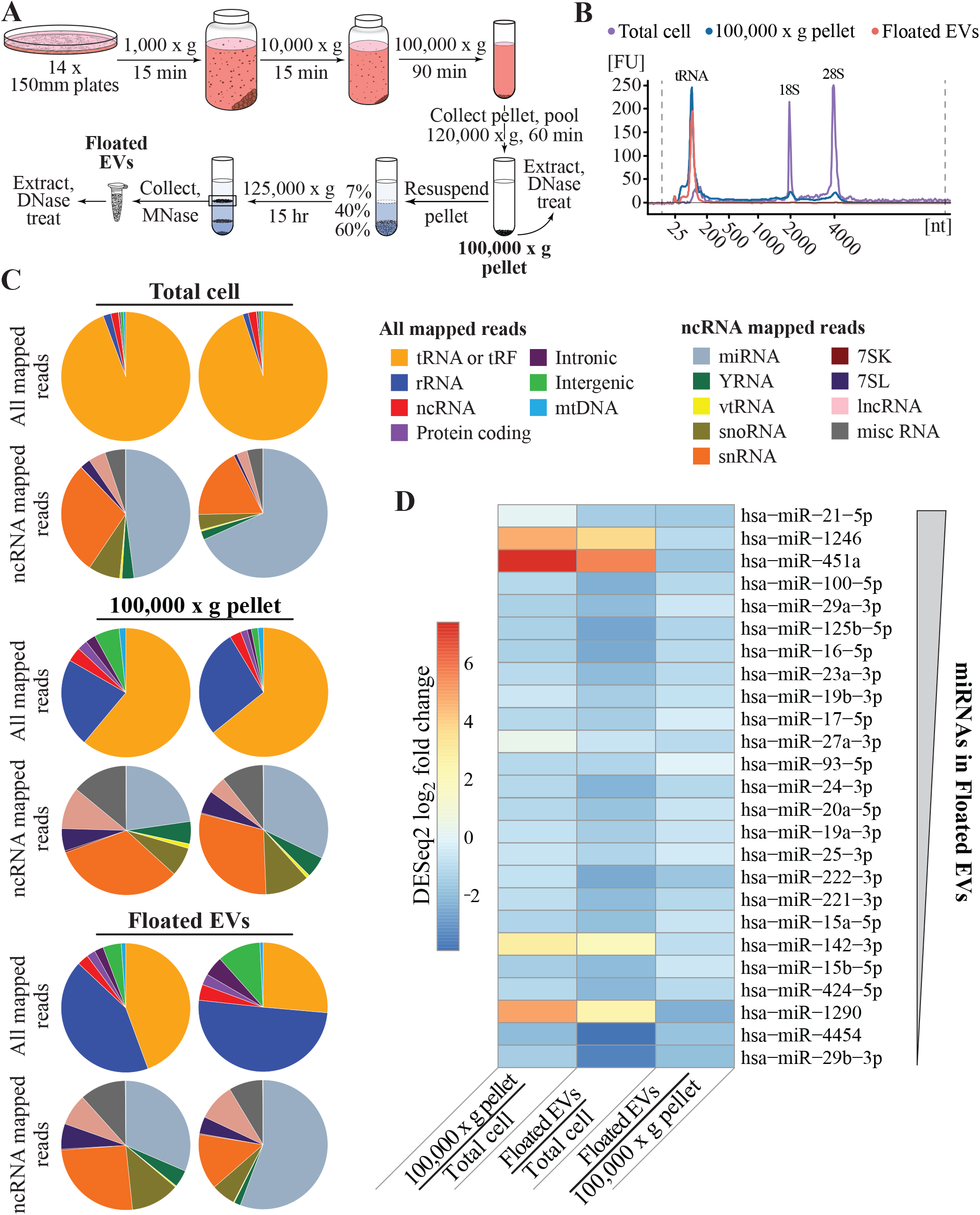
OTTR RNA-seq inventorying of EV sRNA. A) Schematic of EV purification. B) Agilent Bioanalyzer RNA traces for cellular RNA (purple), the 100,000 x g pellet (blue), and floated EVs (peach). Peaks corresponding to tRNA and 18S and 28S rRNA are indicated. C) Pie charts of mapped read assignments of MDA-MB-231 sRNA libraries from 2 biological replicates. tRAX and miRDeep2 were used to map tRNA and miRNA reads, respectively. rRNA, ncRNA, and protein-coding reads aligned to annotated transcripts or genomic ncRNA loci. Intronic, intergenic, and mitochondrial (mt) DNA reads mapped in corresponding locations. Among ncRNA reads, vt is vault and miscRNA includes all ncRNA not split out into other pie slices. D) EV miRNA enrichment in MDA-MB-231 cells based on DESeq2 log_2_ fold change estimates between, as pairwise combinations, 100,000 x g pellet and Total cell, Floated EVs and Total cell, and Floated EVs and 100,000 x g pellet. The 25 miRNAs included in the panel are the most abundant in read count in MDA-MB-231 Floated EVs, with most abundant of the 25 at top as schematized by the thicker end of the gray wedge at right.

Isolated RNA pools were used directly, without gel purification, for OTTR cDNA library synthesis and sequencing. Unsurprisingly, total cellular sRNA and EV library reads were dominated by tRNAs or tRFs and rRNA fragments, as evident from pie charts comparing RNA species across all mapped reads and EV enriched populations (Figs. 5C and S3A, Table S3). Reads from full-length tRNAs as well as tRFs had genome-mismatched nts at expected positions of post-transcriptional modification (Fig. S4).

To evaluate ncRNA representation in more detail, we split the non-tRNA, non-rRNA ncRNA fraction into its own set of pie slices (Figs. 5C and S3A, Table S3). Among ncRNA categories, miRNAs were readily detectable in all samples (grey slices of ncRNA read pies, Figs. 5C and S3A). All samples also contained a sizeable read count from fragments of small nuclear (sn) RNA (orange slices of ncRNA read pies, Figs. 5C and S3A). Additional well-sampled ncRNA in MDA-MB-231 EVs included fragments of long non-coding RNA (lncRNA), small nucleolar RNA (snoRNA), and 7SL RNA (Fig. 5C, Table S3). In HEK 293T cells, 7SL and Y RNA reads were particularly abundant in OTTR libraries from floated EVs (Fig. S3A, Table S3).

Hundreds of miRNA or putative miRNA sequences were detected in EV samples. Considering individual miRNAs ranked in order by their read count in EV libraries, most were as well or better represented in total cellular sRNA than in EVs (Figs. 5D and S3B, note the ratio descriptions at the bottom of the panel). Some exceptions to non-selective EV sorting confirm previous reports of EV-enriched miRNA, for example the relatively abundant miR-451a and miR-142-3p (42, 49, 50). Relative read count abundance of individual miRNAs in total cellular sRNA differed across cell types, but generally EV-enrichment of a particular miRNA did not (compare Figs. 5D and S3B). We identified miRNA enriched in EVs that do not have a bovine counterpart and therefore could not derive from growth of cells in serum prior to their shift to serum-free conditions for EV harvest, such as miR-1290 (Figs. 5D and S3B).

Surprisingly, EV sRNA reads that mapped to some loci annotated as encoding miRNA or lncRNA did not have the consistent 5’ and 3’ end positions expected from cellular processing of functional ncRNA. The relaxed precision of miRNA end-positions in EV samples relative to total cellular sRNA can be visualized in mapped reads for miRNAs including miR-451a (Figs. S3C and S5). We suggest that this heterogeneity is a signature of EV ncRNA, detectable only in NGS libraries that oblige end-to-end RNA sequence capture and do not obscure ncRNA end positions by RNA polynucleotide tailing prior to library synthesis.

## Discussion

Here we describe an approach of controlled serial template jumping to synthesize a cDNA library that captures input sRNA sequences end-to-end. Our approach uses cDNA synthesis to fuse 5′ and 3′ adaptor sequences. Discontinuous templates produce a continuous cDNA with three segments in specific order: the 5′ adaptor primer, the complement of a single template from the input RNA pool, and the 3′ adaptor. To enforce this order of template copying, the primer has a 3′ pyrimidine single-nt overhang, the input pool of templates has a 3′ purine nt, and the 3′ adaptor template has a 3′ cytidine. The comprehensiveness of sRNA sequence capture by OTTR relies on an unanticipated ability of non-retroviral RTs to act as robust terminal transferase enzymes in reactions with Mn^2+^ as the divalent ion. Unique molecular identifier tracts in the 3′ adaptor template can be used to normalize PCR bias, and bar codes are readily added to multiplex libraries. Assay conditions used in this work retained some bias against 3′- labeling of templates with 3′ uridine, but enzyme and buffer modifications can neutralize that bias (HEU, LF, SCP, NTI, and KC, unpublished data).

In addition to the simplicity and low bias of OTTR for NGS library production, the OTTR strategy has additional benefits. First, ddNTP labeling of the input pool 3′ ends precludes templates from the self-priming and cross-priming that creates aberrant cDNA fusions (36, 37). Second, the use of serial template jumps to add flanking adaptors obliges end-to-end copying of input templates. *B. mori* R2 RT can template-jump only when synthesis reaches the 5’ end of an engaged template, as enzyme paused mid-template does not support template jumping (20)(HEU and KC, unpublished data), so partial cDNAs would drop out of the library due to lack of a 3’ adaptor. Third, OTTR requires much less input (as low as 0.2 ng, data not shown) than used in other protocols of precise end-to-end RNA capture, for example in previous TGIRT-seq of similar preparations of EV RNA (42).

It is general consensus in the EV field that better RNA-seq is needed, despite hundreds of millions of dollars invested in EV RNA surveys to date (51–53). The gold-standard 4N method is challenging for non-experts to perform, labor intensive, and not efficient in conversion of input RNA to cDNA. OTTR overcomes all of these barriers. Both reproducibility and conversion efficiency are improved by the limited number of reaction steps during cDNA library preparation, and the simplicity of the protocol makes it amenable to automation. Information gained from sequencing OTTR libraries of EV RNAs adds weight to previous conclusions that EVs contain RNA fragments (14), here established by obligate end-to-end template copying. In addition, the length heterogeneity of EV-enriched sequences from specific miRNA loci suggest the possibility of targeted EV enrichment of misprocessed or misfolded ncRNA, likely combined with EV enrichment of nuclease(s) that fragment structured RNAs (54). The OTTR approach for NGS cDNA library production will be useful for broader inventory and comparison of end-to-end ncRNA sequences present in EVs from human biofluids and tissues in the quest for diagnostic signatures of human health and disease (55).

## Materials and Methods

### Recombinant protein expression and purification

Codon-optimized open reading frames for EuRe and BoMoC RT proteins were ordered from GenScript. Proteins were produced in *Escherichia coli* by expression from MacroLab vector 2bct with a C-terminal 6xHis tag (https://qb3.berkeley.edu/facility/qb3-macrolab/#facility-about) modified to include an N-terminal MBP tag. Cells were grown in 2xYT medium using Rosetta2(DE3)pLysS cells and induced at OD_600_ 0.9 at 16°C overnight with 0.5 mM IPTG. Lysis of the induced cell pellet took place in 20 mM Tris-HCl pH 7.5, 1 M NaCl, 10% glycerol, 1 mM MgCl_2_, 1 mM β-mercaptoethanol (BME), and protease inhibitors by sonication on ice for 3.5 min (10s on, 10s off). Three-step purification was initiated by column binding 6xHis-tagged proteins on nickel agarose. Following binding, the column was washed with 5 volumes of 20 mM Tris HCl pH 7.4, 1 M KCl, 20 mM imidazole, 10% glycerol, and 1 mM BME. Elution from the resin proceeded with 20 mM Tris-HCl pH 7.4, 1M KCl, 400 mM imidazole, 10% glycerol and 1 mM BME. Protein eluted from the Ni column was desalted into heparin buffer A (5 mM HEPES-KOH pH 7.5, 400 mM KCl, 2% glycerol, 0.2 mM dithiothreitol [DTT]) then bound to heparin-Sepharose. The column was washed with 5 column volumes heparin buffer A then ramped up by gradient over 15 column volumes to 20 mM HEPES-KOH pH 7.5, 2 M KCl, 10% glycerol, and 1 mM DTT. Eluted protein peak was pooled and diluted back to 20 mM HEPES-KOH pH 7.5, 400 mM KCl, 10% glycerol, and 1 mM DTT. Protein was size-fractionated using a HiPrep 16/60 Sephacryl S-200HR in 25 mM HEPES-KOH pH 7.5, 0.8 M KCl, 10% glycerol, and 1 mM DTT. Purified protein concentration was determined by UV absorbance using the calculated extinction coefficient (EuRe: 1.325 M^−1^ cm^−1^, BoMoC: 1.350 M^−1^ cm^−1^) and validated as homogeneous full-length RT fusion protein by SDS-PAGE. Final protein concentrations stored at −80°C were 4.1 mg/ml for EuRe and 8.0 mg/ml for BoMoC. Working stock of protein was diluted ~5 fold in 25 mM HEPES-KOH, 0.8 M KCl, 50% glycerol, and 1 mM DTT then moved to −20°C.

### RT assay conditions

Templated cDNA synthesis and NTA assays were carried out under the following conditions: 25 mM Tris-HCl pH 7.5, 75-150 mM KCl, 1-5 mM MgCl_2_, 5 mM DTT, 2.5% glycerol, 90-400 nM RNA-DNA duplex, 45-200 nM RNA or DNA template, 2.5 nM-500 μM dNTP, and 0.5-2 μM enzyme with or without 2% polyethylene glycol (PEG)-6000. Samples were incubated for 30 min to 2 h at 37°C, heat inactivated at 65°C for 5 min, nuclease treated with 0.5 μg/μl RNaseA (Sigma, R6513) and 0.5 units of thermostable RNaseH (NEB, M0523S), then stopped with 10 mM Tris pH 8.0, 0.5 mM ethylenediaminetetraacetic acid (EDTA), and 0.1% sodium dodecyl sulfate (SDS). Products were extracted with phenol:chloroform:isoamyl alcohol (PCI, 25:24:1), ethanol precipitated using 10 μg glycogen as a carrier, and air dried for 5 min prior to resuspending in 5 μl H_2_O. Products were separated by 7.5-15% denaturing urea-PAGE gel then stained using SYBR Gold and imaged by Typhoon Trio.

### Terminal transferase assay conditions

Terminal transferase activity assays were performed under conditions similar to those described above except for the presence of MnCl_2_ rather than MgCl_2_. Briefly, assays contained 25 mM Tris-HCl pH 7.5, 75-150 mM KCl, 5 mM MnCl_2_, 5 mM DTT, 2.5% glycerol, 400 nM RNA-RNA, RNA-DNA, or DNA-DNA duplex or single stranded RNA or DNA template, 500 μM dNTP/NTP/ddNTP, and 0.5 μM enzyme with or without 5% PEG-8000. Samples were incubated for 30 min at 30-37°C, heat inactivated at 65°C for 5 min, and processed as above with or without the post-RT nuclease treatment step.

### Exosome and total cellular sRNA preparation

MDA-MB-231 or HEK 293T cells were grown to ~80% confluency in 14 × 150 mm dishes with 10% exosome-depleted FBS (Exo-FBS from System Biosciences, EXO-FBS-250A-1) in DMEM GlutaMAX media. For purification of bulk EVs, the conditioned media (420 ml) was collected and floating cells and cellular debris were discarded by low- and medium-speed centrifugations (1,000 x g for 15 min and 10,000 x g for 15 min, respectively) at 4°C using a Sorvall R6+ centrifuge with a fixed angle FIBERlite F14-6X500y rotor. The supernatant fraction was spun at 100,000 x g (28,000 RPM) using an SW-28 rotor for 1.5 h. Pellet fractions were resuspended in EV buffer (10 mM HEPES pH 7.4, 0.85% w/v NaCl), pooled and centrifuged at ~120,000 x g (36,000 RPM) in a SW55 rotor for 1 h. Cleared supernatant was discarded and the concentrated high-speed pellet (“100,000 x g pellet”) was lysed directly in 300 μl of TRI reagent (Zymo Research, R2050-1-200). RNA extraction was performed according to instructions for the Direct-zol RNA Miniprep Kit (Zymo Research, R2072). RNA was eluted in 100 μl and treated with the TURBO DNA-free Kit (Thermo, AM1907) and DNase was inactivated according to manufacturer specifications. RNA was then further purified using the RNA Clean & Concentrator Kit (Zymo Research, R1013). RNA was eluted in 13 μl H_2_O; 3 μl was used for Bioanalyzer analysis.

For EV purification via flotation, the pellet collected after the second high-speed spin was resuspended in 200 μl of EV buffer and 2.8 ml of 60% sucrose (in EV buffer) was subsequently added and mixed thoroughly. Aliquots of 40% and 7% sucrose were overlaid sequentially on top and the sample was centrifuged at ~125,000xg (36,500 RPM) in a SW55 rotor for 15 h. The floated fraction corresponding to a mixture of high- and low-density EVs (42) was treated with MNase (NEB, M0247S) to degrade any nucleic acid not contained within the EVs and deactivated with 25 mM ethylene glycol-bis(β-aminoethyl either)-N,N,N′,N′-tetraacetic acid (EGTA) prior to RNA extraction performed as above.

For isolation of total cellular sRNA, a single 150 mm dish of cells was harvested and washed once with PBS. Cells were lysed with 1 ml lysis buffer from the *mir*Vana miRNA Isolation Kit (ThermoFisher, AM1560) and RNA extraction was performed as specified by the manufacturer. RNA was eluted in 100 μl and treated as above to remove contaminating DNAs and concentrate the sample. Final elution of sRNAs was in 25 μl H_2_O; 4 μl was used for NanoDrop and Bioanalyzer analysis.

### OTTR library generation and sequencing

Input RNA (10 ng) (miRXplore Universal Reference Standard [Miltenyi Biotech, 130-094-407], EV sRNA, or *mir*Vana size-selected total cellular RNA) was diluted into 20 mM Tris-HCl pH 7.5, 150 mM KCl, 0.5 mM DTT, 5% PEG-8000, 2 mM MnCl_2_, 250 μM ddATP (+/− 250 μM ddGTP), and 0.7 μM BoMoC then incubated for 1.5-2 h at 30°C. For ddGTP chase of initial labeling with ddATP, the reaction was allowed to proceed for 1.5 h at 30°C with ddATP only, then chased with 250 μM ddGTP and incubated for another 30 min at 30°C. The reaction was stopped by incubating at 65°C for 5 min followed by addition of 5 mM MgCl_2_ and 0.5 units of Shrimp Alkaline Phosphatase (rSAP, NEB M0371S). The phosphatase reaction was incubated at 37°C for 15 min, stopped by addition of 5 mM EGTA, then incubated at 65°C for 5 min. Subsequently, buffers were added to give an additional 0.5 mM MgCl_2_ and 45 mM KCl plus 2% PEG-6000, 200 μM dGTP, 40 μM dTTP and dCTP, 2 μM dATP +/− 150 μM 2-amino-2′-deoxyadenosine-5′-triphosphate (DAP), 90 nM RNA-DNA primer-duplex with +1T and +1G overhangs, 180 nM terminating AT, and 0.5 μM BoMoC. Samples were processed as above then resuspended in 10 μl H_2_O. If Universal adaptors were used in the RT reaction, libraries were generated using 5 μl of the purified RT reaction product and 4-8 cycles of PCR with Q5 high fidelity polymerase (NEB, M0491S). PCR reaction products were column purified, separated by 7.5% urea-PAGE to remove adaptor dimer, and the desired product was isolated by diffusion overnight at 37°C followed by PCI extraction, ethanol precipitation, and resuspension in 10 μl H_2_O. If Full-length adaptors were used, the RT reaction was treated with RNaseA and RNaseH prior to clean-up and no PCR amplification was performed. Quantification of libraries prior to sequencing used qPCR with primers specific to the Illumina P5 and P7 adaptor sequences and standards from the NEBNext Library Quant Kit (NEB, E7630S). Sequencing of prepared libraries was performed using an Illumina MiniSeq with the 75-cycle high-output kit. Library yield using Universal adaptors ranged from 20-30 nM following 4 cycles of PCR the miRXplore reference standard and from 6-10 nM with more complex input pools (total cellular sRNA and EV RNA) following 4 cycles of PCR. Yield using Full-length adaptors with the miRXplore reference standard was ~ 1 nM. To recover 10M reads for total cellular and EV RNA sequencing, ~1.5% of each library was taken to produce the starting 1 nM sample pool.

### miRXplore benchmarking

SRA accession numbers are given in Table S2 for paired replicates representing QIAseq, TruSeq, CleanTag, NEXTflex, NEBNext, 4N, TGIRT, and SMARTer RNA-seq (35, 38–40). Reads were downloaded using ***fastq-dump***. Adaptors were trimmed using ***cutadapt*** (56) based on manufactures suggestions. All libraries were aligned to a reference of 962 miRXplore miRNAs (Supplementary Dataset S1) using ***bowtie*** (57) with the following parameters: ***--norc -v 0 -m 10 --tryhard --best --strata***. Alignment files raw counts were gathered from the alignments before being normalized by ***DESeq2*** (58) then to counts per million (CPM). Coefficients of variation were measured from the CPM, and a miRNA was considered detectable if it had a CPM > 2. Δlog_2_(CPM) for each miRNA was determined by Δlog_2_(CPM) = log_2_(CPM_observed_)−log_2_(CPM_expected_), where CPM_expected_ is the expected counts from a truly equimolar sampling of the miRXplore miRNAs. ***Pheatmap*** R package was used to compute a correlation distance matrix of the Δlog_2_(CPM) for underweighted pair group method with arithmetic mean (UPGMA) hierarchical cluster (Fig. 4A).

### Bias analysis of miRNA 5′ and 3′ termini

A random forest regression was trained on the three 5′-most and three 3′-most nucleotides as variables to predict the computed Δlog_2_(CPM) of each miRNA, as previously done (40). The ***randomForest*** R package was used to generate 250 random trees with two variables randomly sampled as candidates at each split. Increase in mean square error for each variable (the three 5′-most and three 3′-most nucleotides for each miRNA) (Figs. 4C and 4D) was investigated along with percent variance explained by the model (Fig. 4B) to evaluate importance of miRNA terminal sequence in predicting over- or under-representation in a cDNA library.

### Total, EV pellet, and floated EV RNA read analysis

OTTR oligo sequences were trimmed from the libraries using ***cutadapt*** and reads shorter than 10 bases were discarded. tRNA reads were determined and quantified by ***tRAX*** (59) first, following miRNAs detection and quantification by ***miRDeep2*** (60). The remaining reads greater than 20 bases were retained, and sequentially mapped to human rRNA (**U13369.1, NR_145819.1, NR_146144.1, NR_146151.1, NR_146117.1, X12811.1, ENST00000389680.2, ENST00000387347.2, NR_003287.4, NR_023379.1, NR_003285.3, NR_003286.4**), ncRNA (***Ensembl***), mRNA (***GENCODE***), lncRNA (***GENCODE***), and gDNA (***GENCODE***) using ***bowtie***. The alignment files were merged and ***RSEM*** (61) was used to quantify readcounts. ***Tximport*** and ***DEseq2*** were used to import the counts data and estimate difference expression. Given the wide variation in library composition, analysis was largely restricted to differential expression of miRNA and tRNA.

## Competing interests statement

HEU, LF, SCP, and KC are named inventors on UC Berkeley patent applications describing OTTR technology. HEU and KC are founders of Karnateq Inc., which licensed this technology.

## Author contributions

HEU, LF, NTI, and KC designed research; MMT and XL purified EV; LF performed bioinformatic analyses, SCP contributed to OTTR design and assay development, and HEU performed all other research. HEU, LF, MMT, NTI, and KC analyzed data. HEU, LF, and KC wrote the paper with input from all authors.

## Acknowledgements

HEU, LF, SCP and KC were supported by NIH R35 GM130315 and DP1 HL156819 as well as the Bakar Fellows Program (HEU) and NIH T32 GM007232 (LF). MMT, XL, and RS were supported by Howard Hughes Medical Institute.

---

**Table S1.**
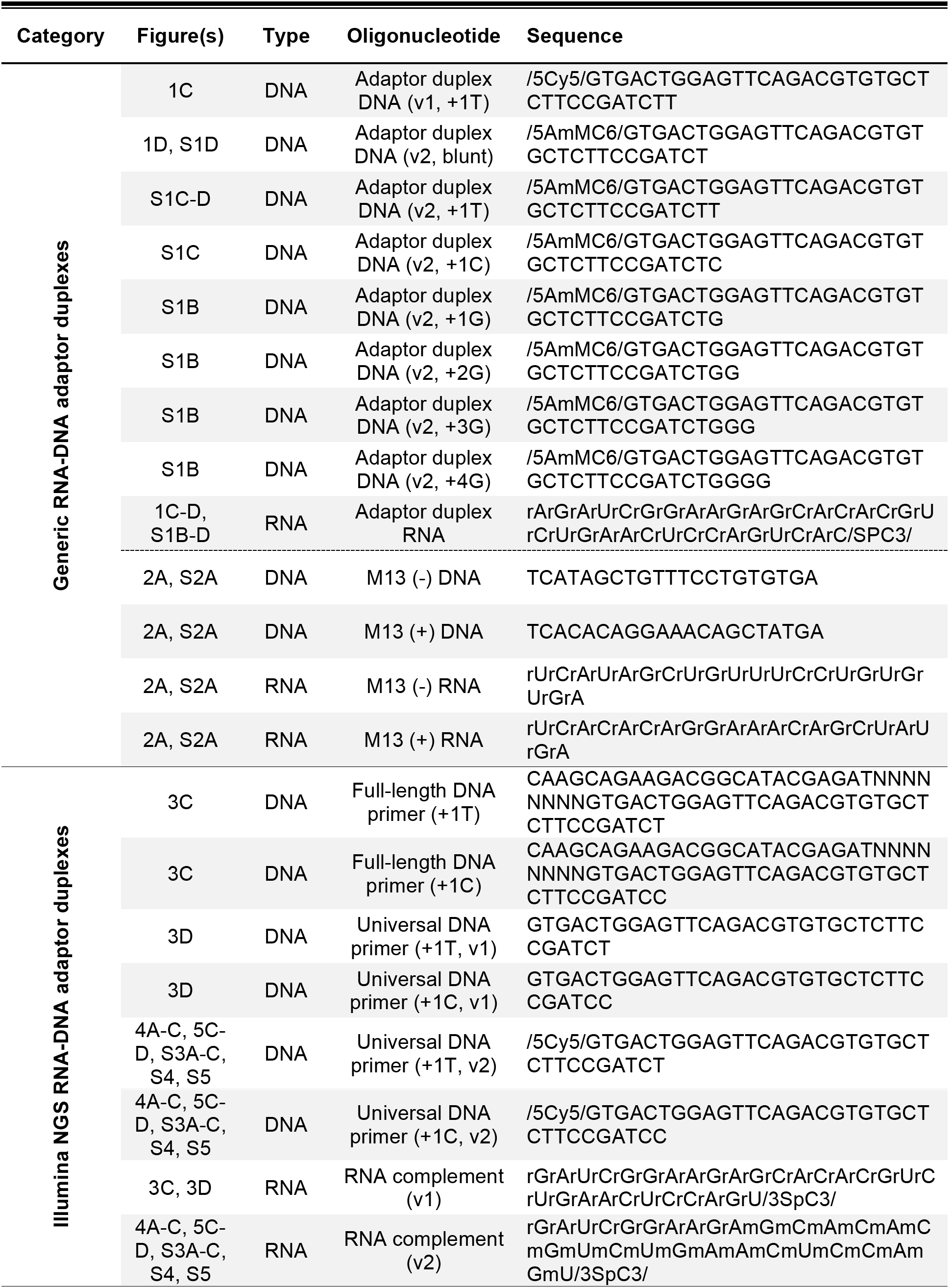
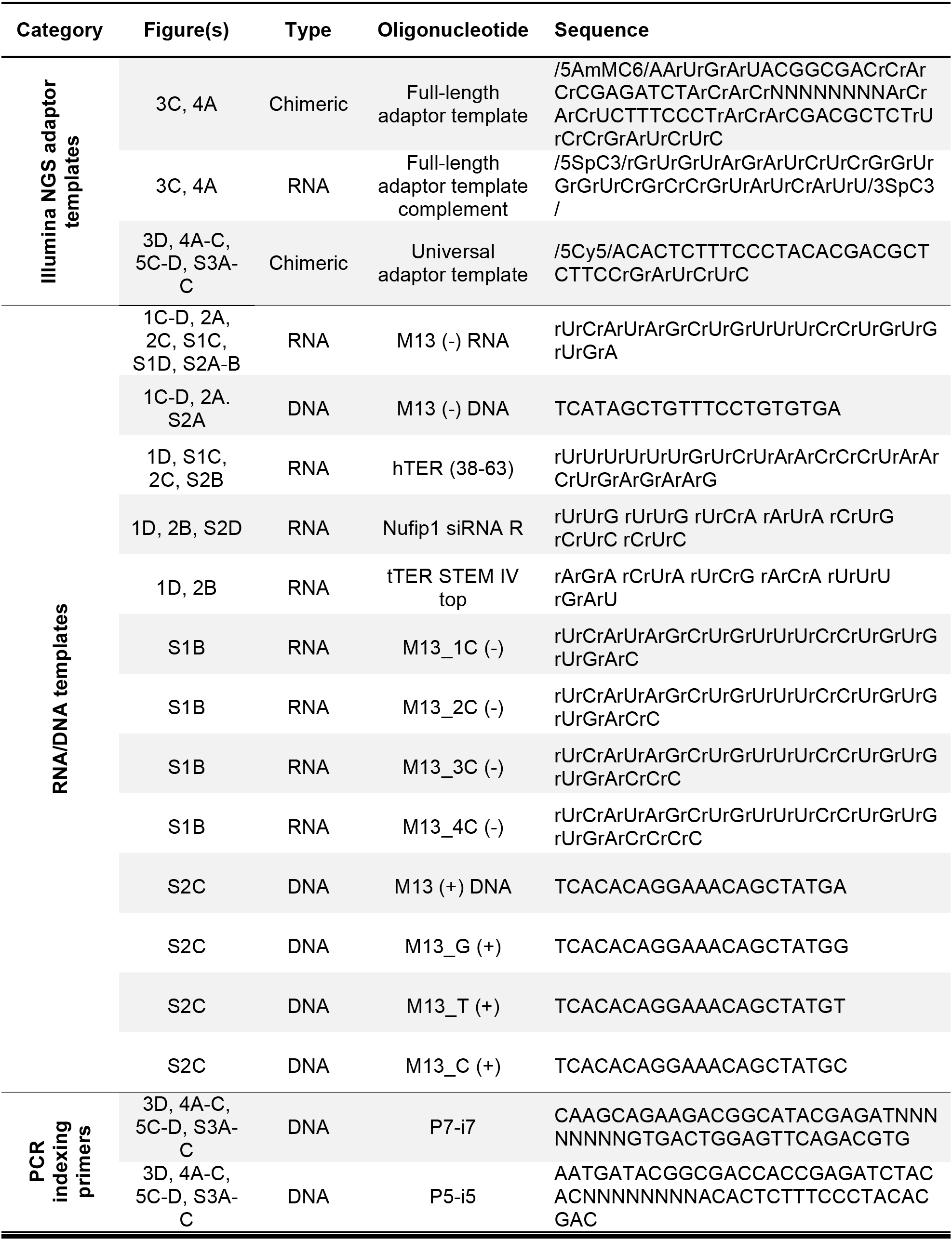
Oligonucleotide sequences used in this study.

**Table S2.** RNA-seq read data for miRXplore miRNA library analysis.

| **Library** | **Manufacturer** | **Replicate Name** | **SRA Number** | **Citation** |
| --- | --- | --- | --- | --- |
| 4N Protocol B | In-house | 4N_B.Lab5.1 | SRR6380698 | 36 |
|  |  | 4N_B.Lab5.2 | SRR6380697 |  |
| NEBNext Small RNA Library Prep Set | New England Biolabs | NEB1 | SRR7777351 | 37 |
|  |  | NEB2 | SRR7777350 |  |
| QIAseq miRNA Library Kit | Qiagen | QIA1 | SRR7777353 | 37 |
|  |  | QIA2 | SRR7777352 |  |
| Nextflex Small RNA-Seq Kit v3 | Bioo Scientific | BSC1 | SRR7777357 | 37 |
|  |  | BSC2 | SRR7777356 |  |
| SMARTer smRNA-Seq Kit | Clontech/Takara | CLT1 | SRR7777363 | 37 |
|  |  | CLT2 | SRR7777362 |  |
| TruSeq Small RNA Library Preparation Kit | Illumina | TruSeq1 | SRR5234319 | 38 |
|  |  | TruSeq2 | SRR5234320 |  |
| CleanTag Small RNA Library Preparation Kit | TriLink BioTechnologies | CleanTag1 | SRR5234287 | 38 |
|  |  | CleanTag2 | SRR5234288 |  |
| Thermostable group II intron reverse transcriptase sequencing | InGex | TGIRT-NTT1 | SRR8186015 | 39 |
|  |  | TGIRT-NTT2 | SRR8186014 |  |

**Table S3.** Percentage of reads mapped to different RNA classes used to generate pie graphs.

| RNA population | MDA-MB-231 |  |  |  |  |  | HEK 293T |  |  |  |
| --- | --- | --- | --- | --- | --- | --- | --- | --- | --- | --- |
|  | Total cell |  | 100,000 x g pellet |  | Floated EVs |  | Total cell |  | Floated EVs |  |
|  | Replicate 1 | Replicate 2 | Replicate 1 | Replicate 2 | Replicate 1 | Replicate 2 | Replicate 1 | Replicate 2 | Replicate 1 | Replicate 2 |
| tRNA or tRF | 94.31% | 94.87% | 61.11% | 64.21% | 44.41% | 26.39% | 72.62% | 79.29% | 87.33% | 67.23% |
| rRNA | 0.14% | 0.09% | 3.05% | 2.47% | 4.96% | 4.61% | 0.14% | 0.17% | 0.13% | 0.04% |
| mt-rRNA | 1.82% | 1.31% | 19.16% | 24.68% | 37.63% | 45.72% | 5.50% | 5.03% | 9.25% | 30.80% |
| protein coding | 0.12% | 0.11% | 2.55% | 1.70% | 2.21% | 2.75% | 1.59% | 0.94% | 0.35% | 0.22% |
| intronic | 0.37% | 0.39% | 2.55% | 1.03% | 2.27% | 4.98% | 4.92% | 3.09% | 0.54% | 0.29% |
| intergenic | 0.77% | 0.74% | 6.28% | 1.66% | 4.58% | 10.80% | 10.46% | 6.76% | 0.98% | 0.50% |
| mtDNA | 0.58% | 0.44% | 1.69% | 1.42% | 1.13% | 0.80% | 0.43% | 0.73% | 0.05% | 0.02% |
| miRNA | 0.91% | 1.39% | 0.82% | 0.91% | 0.89% | 2.20% | 3.11% | 2.62% | 0.64% | 0.23% |
| YRNA | 0.06% | 0.04% | 0.20% | 0.15% | 0.12% | 0.06% | 0.03% | 0.05% | 0.12% | 0.13% |
| vtRNA | 0.01% | 0.01% | 0.04% | 0.02% | 0.01% | 0.01% | 0.02% | 0.03% | 0.01% | 0.01% |
| snoRNA | 0.15% | 0.08% | 0.26% | 0.31% | 0.34% | 0.24% | 0.09% | 0.22% | 0.02% | 0.03% |
| snRNA | 0.54% | 0.36% | 1.18% | 0.84% | 0.72% | 0.55% | 0.15% | 0.40% | 0.13% | 0.15% |
| 7SK | 0.00% | 0.00% | 0.02% | 0.01% | 0.01% | 0.01% | 0.01% | 0.00% | 0.00% | 0.00% |
| 7SL | 0.05% | 0.02% | 0.20% | 0.16% | 0.18% | 0.17% | 0.17% | 0.09% | 0.18% | 0.12% |
| lncRNA | 0.08% | 0.05% | 0.38% | 0.13% | 0.22% | 0.37% | 0.47% | 0.38% | 0.08% | 0.06% |
| misc RNA | 0.10% | 0.08% | 0.50% | 0.30% | 0.33% | 0.34% | 0.30% | 0.19% | 0.19% | 0.18% |

**Figure S1.**
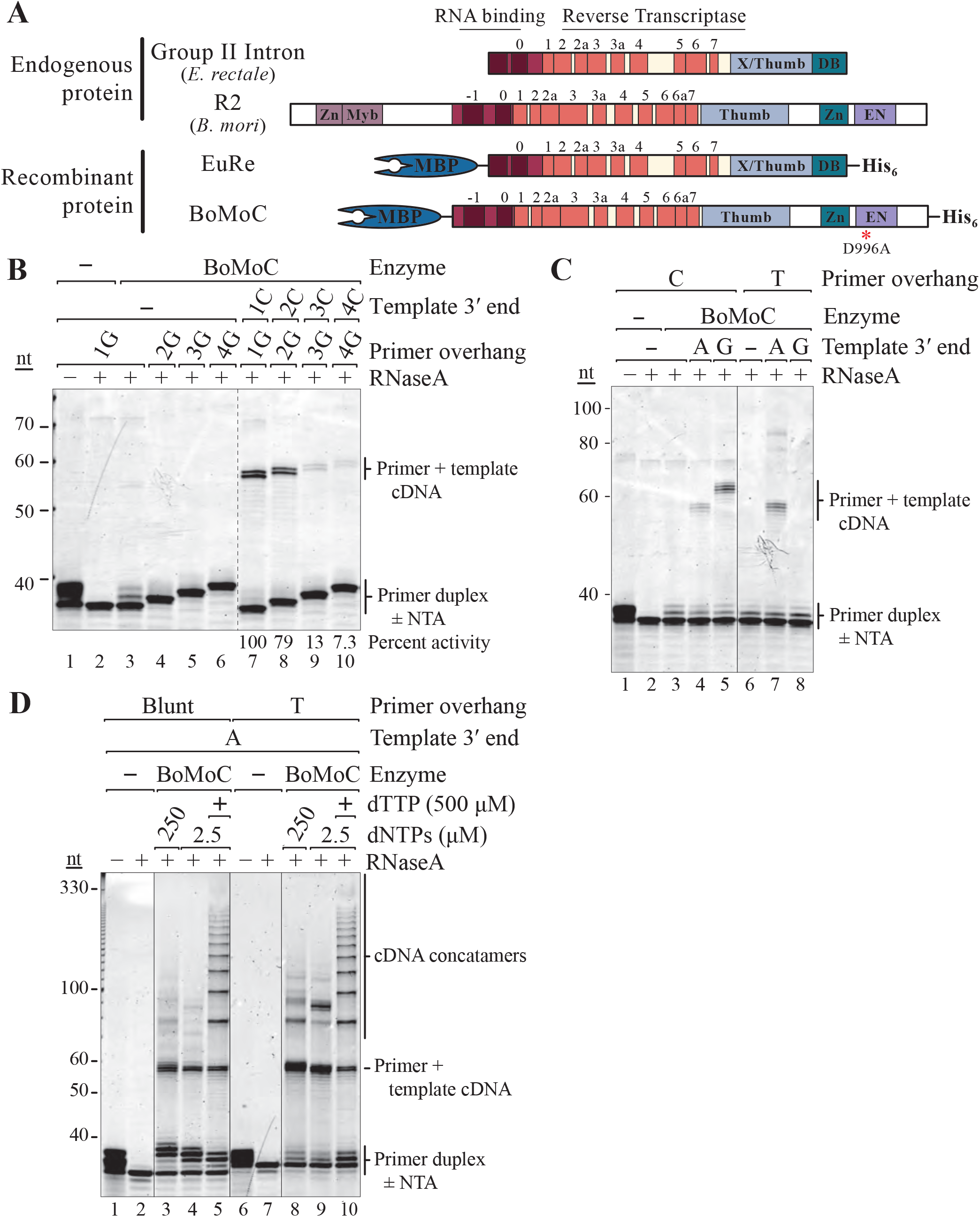
Primer requirements for template jumping by BoMoC. A) Domain layout for the endogenous and modified RTs used in this work. The group II intron RT from *E. rectale* and the non-LTR R2 RT from *B. mori* (top) were used to generate tagged proteins for purification and characterization (bottom). Numbering is for RT active site motifs, and other domains are described in the main text. B, C, D) SYBR Gold-stained denaturing PAGE gel showing the activity of BoMoC on DNA-RNA primer duplexes. In (B), The DNA primer had +1G, +2G, +3G, or +4G overhang in the absence (lanes 3-6) or presence (lanes 7-10) of RNA template with a 3’ end complementary to the duplex overhang. Normalized cDNA synthesis activity was quantified as indicated below the lanes. In (C), a +1C (lanes 3-5) or +1T (lanes 6-8) primer was used with RNA templates with 3’ A or 3’ G end. In (D), a blunt-end (lanes 3-5) or +1T overhang (lanes 8-10) primer was assayed with 3’A RNA template in the presence of different concentrations of mixed dNTPs and extra dTTP.

**Figure S2.**
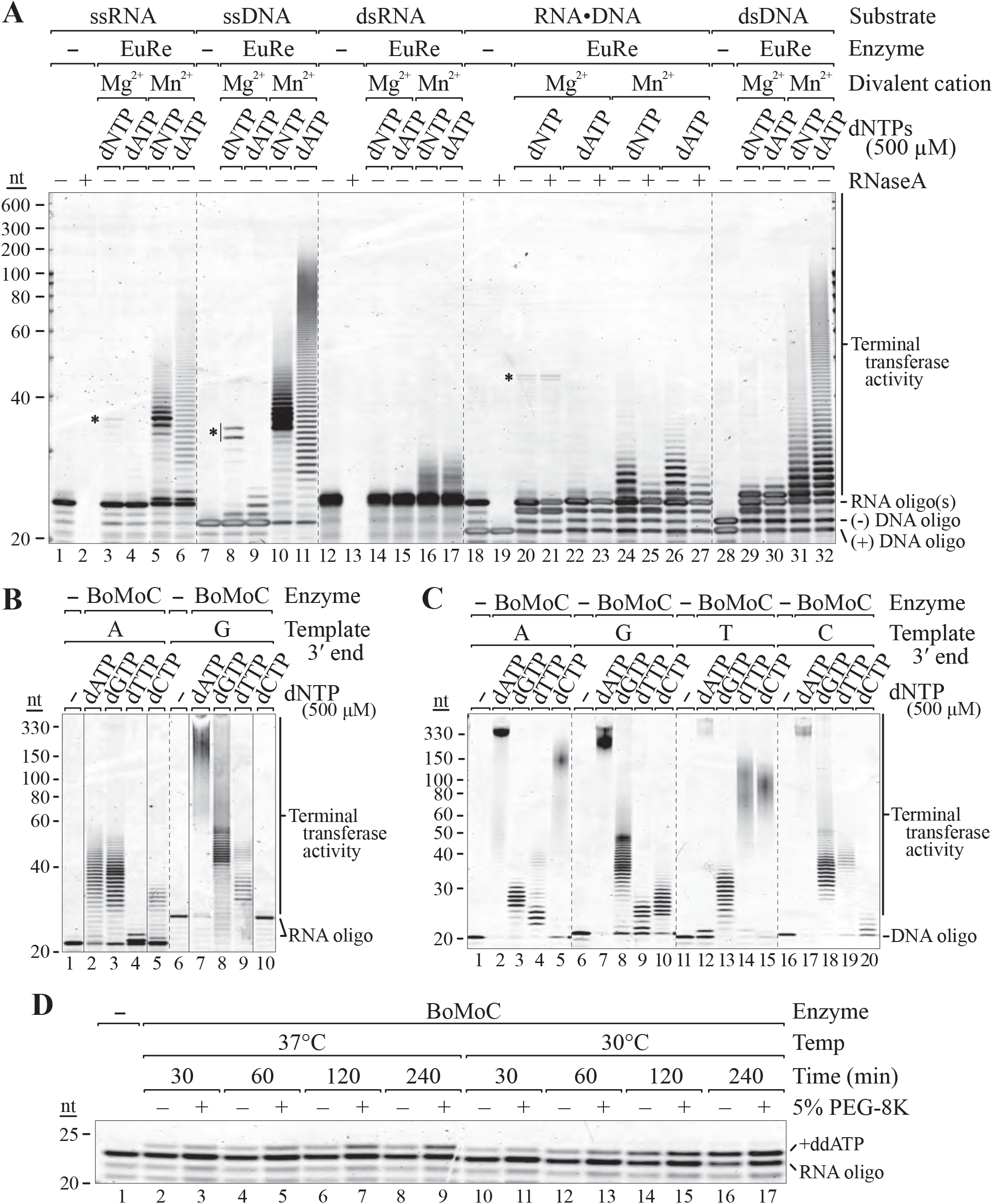
Terminal transferase activity of EuRe and BoMoC RTs. SYBR Gold-stained denaturing PAGE gels showing A) EuRe activity on ssRNA or ssDNA or blunt-ended dsRNA, dsDNA, or RNA-DNA duplex under Mg^2+^ or Mn^2+^ conditions in the presence of 500 uM dNTP or dATP. Products marked with asterisks indicate non-complementary oligonucleotide cross-priming in reactions with Mg^2+^. B) BoMoC activity on ssRNA templates with 3’ A and 3’ G in Mn^2+^ conditions in the presence of individual dNTPs. C) BoMoC activity on ssDNA templates with 3’ A, G, T, or C under Mn^2+^ conditions in the presence of individual dNTPs. D) BoMoC terminal transferase activity in Mn^2+^ using an inefficiently tailed 3’ U RNA, with or without 5% PEG-8K at 37°C or 30°C for 30 min to 4 h.

**Figure S3.**
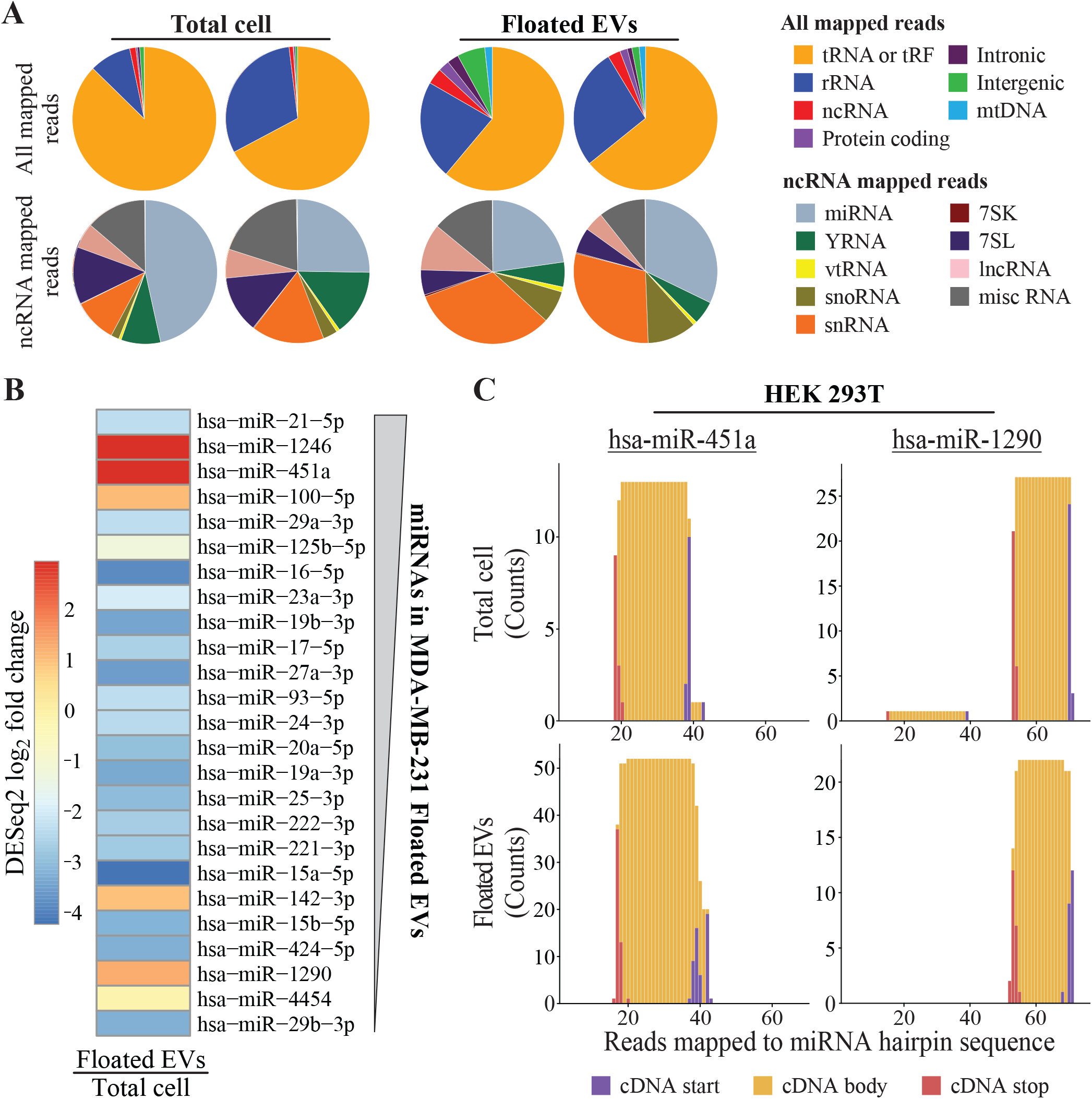
Analysis of OTTR cDNA libraries from HEK 293T-cell RNA pools. A) Pie charts of mapped reads from HEK 293T OTTR sRNA libraries. See Figure 5 legend for definitions. B) EV miRNA enrichment in HEK 293T cells based on DESeq2 log_2_ fold change estimates between Floated EVs and Total cell. The 25 miRNAs included in the heat map are the most abundant in read count in MDA-MB-231 Floated EVs for comparison to Figure 5D; the most abundant of the 25 is at top, as schematized by the thicker end of the gray wedge. C) Cumulative number of aligned bases across loci encoding hsa-miR-451a and hsa-miR-1290. Purple marks the position of the RNA 3’ end/ cDNA5’ end and red marks the position of the RNA 5’ end/ cDNA 3’ end based on end-to-end sequence capture. Y-axes are unnormalized mapping read counts from total mapped read counts of 4,048,198 and 2,190,523 for Total cell and Floated EVs, respectively.

**Figure S4.**
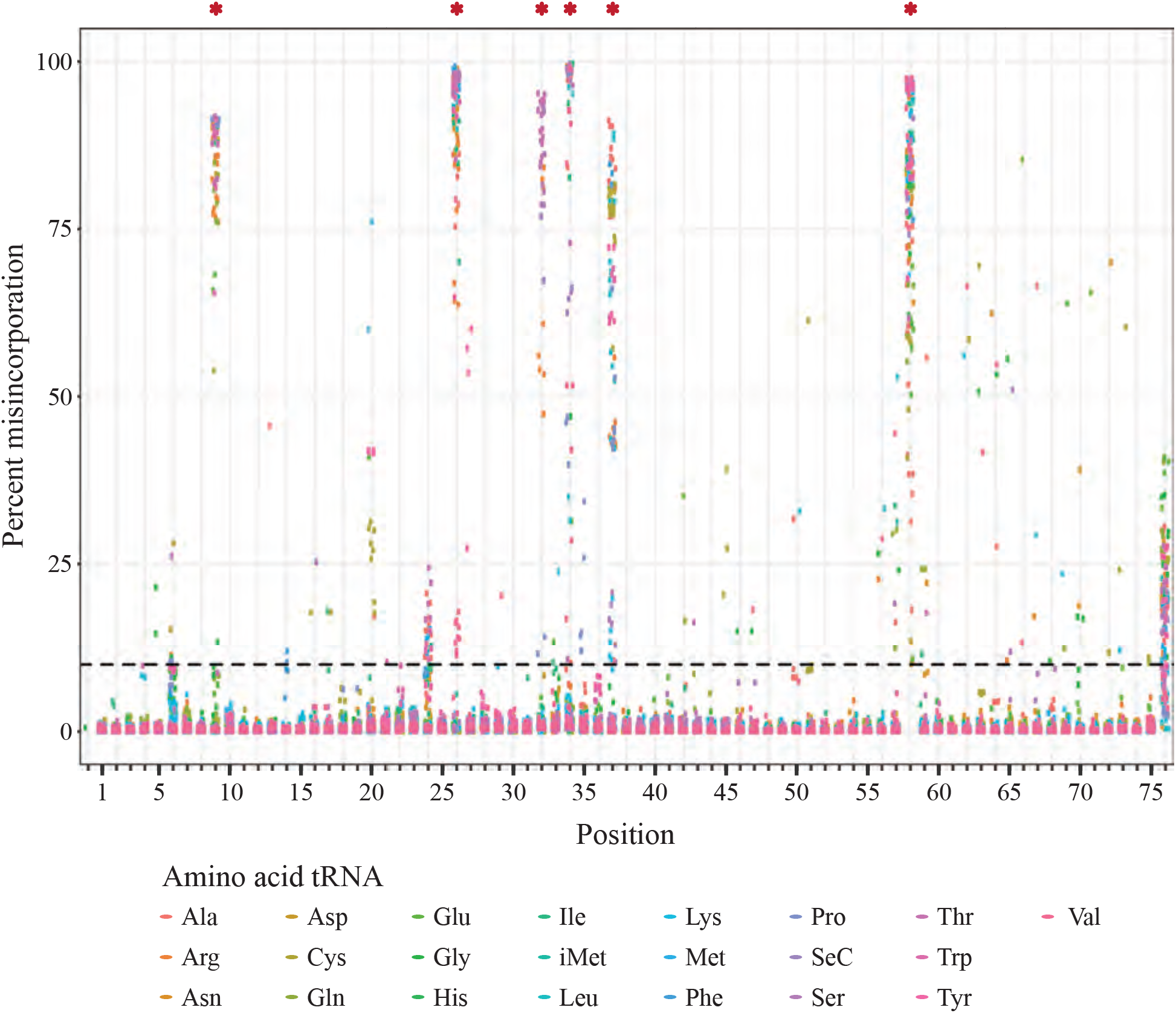
Frequency of discordance between tRNA reads and genome sequence for MDA-MB-231 total cellular sRNA. Asterisks indicate general positions of tRNA post-transcriptional base modification at positions 9 (*N*^1^-methylguanosine and *N*^1^-methyladenosine), 26 (*N*^2^-methylguanosine and *N*^2^,*N*^2^-dimethylguanosine), 32 (2′-O-methylcytidine, 2′-O-methyluridine, 2′- O-methylpseudouridine, pseudouridine, and 3-methylcytidine), 34 (pseudouridine, 5-carbamoylmethyluridine, 5-methoxycarbonylmethyluridine, 5-(carboxyhydroxymethyl)uridine methyl ester, 5-methoxycarbonylmethyl-2′-O-methyluridine, 5-methoxycarbonylmethyl-2-thiouridine, inosine, 5-taurinomethyluridine, 5-carboxymethylaminomethyluridine, 5-taurinomethyl-2-thiouridine, 2′-O-methylcytidine, 5-hydroxymethylcytidine, 2′‐O‐Methyl-5-hydroxymethylcytidine, 5-formyl-2′-O-methylcytidine, 2′-O-methylguanosine, queuosine, mannosyl-queuosine, galactosyl-queuosine, 5-carboxymethylaminomethyl-2-thiouridine, and 5-formylcytidine), 37 (*N*^1^-methylguanosine, *N*^6^-isopentenyladenosine, <u>*N*^6^</u>- <u>threonylcarbamoyladenosine</u>, <u>*N*^6^</u>-methyl-<u>*N*^6^</u>-threonylcarbamoyladenosine, 1-methylinosine, wybutosine, 2-methylthio-*N*^6^-threonylcarbamoyladenosine, hydroxywybutosine, and <u>peroxywybutosine</u>), and 58 (1-methyladenosine). See Suzuki, T. The expanding world of tRNA modifications and their disease relevance. *Nat Rev Mol Cell Biol* (2021).

**Figure S5.**
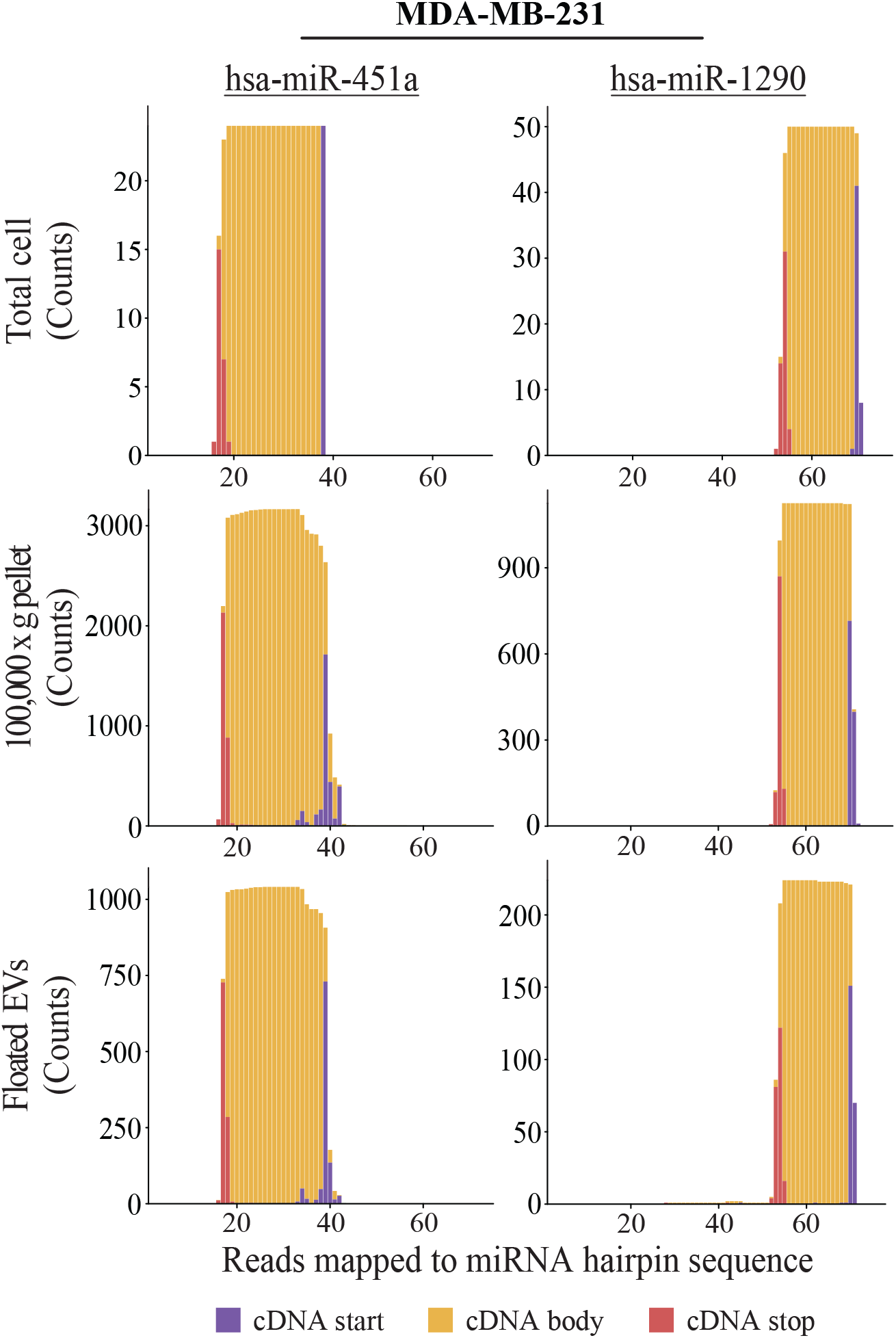
Read plots for miR-451a and miR-1290 in MDA-MB-231 OTTR RNA-seq libraries. Cumulative number of aligned bases across loci encoding hsa-miR-451a and hsa-miR-1290 from each OTTR sRNA library is shown. Purple marks the position of the RNA 3’ end/ cDNA5’ end and red marks the position of the RNA 5’ end/ cDNA 3’ end based on end-to-end sequence capture. Y-axes are unnormalized mapping read counts from total mapped read counts of 14,341,167, 4,405,336, and 1,098,314 for Total cell, 100,000 x g pellet, and Floated EVs, respectively.

## References

1. Lander ES, et al. (2001) Initial sequencing and analysis of the human genome. Nature 409:860–921.

2. Eickbush TH & Jamburuthugoda VK (2008) The diversity of retrotransposons and the properties of their reverse transcriptases. Virus Res 134:221–234.

3. Sultana T, Zamborlini A, Cristofari G, & Lesage P (2017) Integration site selection by retroviruses and transposable elements in eukaryotes. Nat Rev Genet 18:292–308.

4. Eickbush TH & Eickbush DG (2015) Integration, Regulation, and Long-Term Stability of R2 Retrotransposons. Microbiol Spect 3:Mdna3-0011–2014.

5. Fujiwara H (2015) Site-specific non-LTR retrotransposons. Microbiol Spect 3:Mdna3-0001–2014.

6. Luan DD & Eickbush TH (1995) RNA template requirements for target DNA-primed reverse transcription by the R2 retrotransposable element. Mol Cell Biol 15:3882–3891.

7. Lambowitz AM & Belfort M (2015) Mobile bacterial group II introns at the crux of eukaryotic evolution. Microbiol Spect 3:MDNA3-0050–2014.

8. Zhu YY, Machleder EM, Chenchik A, Li R & Siebert PD (2001) Reverse transcriptase template switching: a SMART approach for full-length cDNA library construction. BioTechniques 30: 892–897.

9. Wulf MG, et al. (2019) Non-templated addition and template switching by Moloney murine leukemia virus (MMLV)-based reverse transcriptases co-occur and compete with each other. J Biol Chem 294:18220–18231.

10. Negroni M & Buc H (2001) Retroviral recombination: what drives the switch? Nat Rev Mol Cell Biol 2:151–155.

11. Boivin V, et al. (2020) Reducing the structure bias of RNA-Seq reveals a large number of non-annotated non-coding RNA. Nucl Acids Res 48:2271–2286.

12. Ozsolak F & Milos PM (2011) RNA sequencing: advances, challenges and opportunities. Nat Rev Genet 12:87–98.

13. Stark R, Grzelak M, & Hadfield J (2019) RNA sequencing: the teenage years. Nat Rev Genet 20:631–656.

14. Abramowicz A & Story MD (2020) The long and short of it: The emerging roles of non-coding RNA in small extracellular vesicles. Cancers 12:1445.

15. Guo LT, et al. (2020) Sequencing and structure probing of long RNAs using MarathonRT: A next-generation reverse transcriptase. J Mol Biol 432:3338–3352.

16. Zhao C, Liu F, & Pyle AM (2018) An ultraprocessive, accurate reverse transcriptase encoded by a metazoan group II intron. RNA 24:183–195.

17. Katibah GE, et al. (2014) Broad and adaptable RNA structure recognition by the human interferon-induced tetratricopeptide repeat protein IFIT5. Proc Natl Acad Sci USA 111:12025–12030.

18. Yang J, Malik HS, & Eickbush TH (1999) Identification of the endonuclease domain encoded by R2 and other site-specific, non-long terminal repeat retrotransposable elements. Proc Natl Acad Sci USA 96:7847–7852.

19. Zhao C & Pyle AM (2016) Crystal structures of a group II intron maturase reveal a missing link in spliceosome evolution. Nat Struct Mol Biol 23:558–565.

20. Bibillo A & Eickbush TH (2004) End-to-end template jumping by the reverse transcriptase encoded by the R2 retrotransposon. J Biol Chem 279:14945–14953.

21. Oz-Gleenberg I, Herschhorn A, & Hizi A (2011) Reverse transcriptases can clamp together nucleic acids strands with two complementary bases at their 3′-termini for initiating DNA synthesis. Nucl Acids Res 39:1042–1053.

22. Clark JM, Joyce CM, & Beardsley GP (1987) Novel blunt-end addition reactions catalyzed by DNA polymerase I of Escherichia coli. J Mol Biol 198:123–127.

23. Holton TA & Graham MW (1991) A simple and efficient method for direct cloning of PCR products using ddT-tailed vectors. Nucl Acids Res 19:1156.

24. Marchuk D, Drumm M, Saulino A, & Collins FS (1991) Construction of T-vectors, a rapid and general system for direct cloning of unmodified PCR products. Nucl Acids Res 19:1154–1154.

25. Clark JM (1988) Novel non-templated nucleotide addition reactions catalyzed by procaryotic and eucaryotic DNA polymerases. Nucl Acids Res 16:9677–9686.

26. Golinelli M-P & Hughes SH (2002) Nontemplated base addition by HIV-1 RT can induce nonspecific strand transfer in vitro. Virology 294:122–134.

27. Patel PH & Preston BD (1994) Marked infidelity of human immunodeficiency virus type 1 reverse transcriptase at RNA and DNA template ends. Proc Natl Acad Sci USA 91:549–553.

28. Ohtsubo Y, Nagata Y & Tsuda M (2017) Efficient N-tailing of blunt DNA ends by Moloney murine leukemia virus reverse transcriptase. Sci Rep 7: 41769.

29. Luczkowiak J, Matamoros T & Menéndez-Arias L (2018) Template-primer binding affinity and RNase H cleavage specificity contribute to the strand transfer efficiency of HIV-1 reverse transcriptase. J Biol Chem 293:13351–13363.

30. Martin G & Keller W (2007) RNA-specific ribonucleotidyl transferases. RNA 13:1834–1849.

31. Motea EA & Berdis AJ (2010) Terminal deoxynucleotidyl transferase: The story of a misguided DNA polymerase. Biochim Biophys Acta 1804:1151–1166.

32. Bibiłło A & Eickbush TH (2002) The reverse transcriptase of the R2 non-LTR retrotransposon: continuous synthesis of cDNA on non-continuous RNA templates. J Mol Biol 316:459–473.

33. Oz-Gleenberg I, Herzig E, & Hizi A (2012) Template-independent DNA synthesis activity associated with the reverse transcriptase of the long terminal repeat retrotransposon Tf1. FEBS 279:142–153.

34. Brown JA & Suo Z (2011) Unlocking the sugar “steric gate” of DNA polymerases. Biochemistry 50:1135–1142.

35. Giraldez MD, et al. (2019) Comprehensive multi-center assessment of small RNA-seq methods for quantitative miRNA profiling. Nat Biotechnol 36:746–757.

36. Kaushik N, et al. (2000) Role of glutamine 151 of human immunodeficiency virus type-1 reverse transcriptase in substrate selection as assessed by site-directed mutagenesis. Biochemistry 39:2912–2920.

37. Bibillo A & Eickbush TH (2002) High processivity of the reverse transcriptase from a non-long terminal repeat retrotransposon. J Biol Chem 277:34836–34845.

38. Coenen-Stass AML, et al. (2018) Evaluation of methodologies for microRNA biomarker detection by next generation sequencing. RNA Biol 15:1133–1145.

39. Shore S, et al. (2016) Small RNA library preparation method for next-generation sequencing using chemical modifications to prevent adapter dimer formation. PLoS One 11:e0167009.

40. Xu H, Yao J, Wu DC, & Lambowitz AM (2019) Improved TGIRT-seq methods for comprehensive transcriptome profiling with decreased adapter dimer formation and bias correction. Sci Rep 9:7953.

41. Mohr S, et al. (2013) Thermostable group II intron reverse transcriptase fusion proteins and their use in cDNA synthesis and next-generation RNA sequencing. RNA 19:958–970.

42. Temoche-Diaz MM, et al. (2019) Distinct mechanisms of microRNA sorting into cancer cell-derived extracellular vesicle subtypes. eLife 8:e47544.

43. Everaert C, et al. (2019) Performance assessment of total RNA sequencing of human biofluids and extracellular vesicles. Sci Rep 9:17574.

44. Morris KV & Mattick JS (2014) The rise of regulatory RNA. Nat Rev Genet 15:423–437.

45. Krishna S, Raghavan S, DasGupta R, & Palakodeti D (2021) tRNA-derived fragments (tRFs): establishing their turf in post-transcriptional gene regulation. Cell Mol Life Sci 78:2607–2619.

46. Lin C-P & He L (2017) Noncoding RNAs in Cancer Development. Ann Rev Cancer Biol 1:163–184.

47. Hulstaert E, et al. (2020) Charting Extracellular Transcriptomes in The Human Biofluid RNA Atlas. Cell Rep 33:108552.

48. Shurtleff MJ, Temoche-Diaz MM, & Schekman R (2018) Extracellular Vesicles and Cancer: Caveat Lector. Annu Rev Cancer Biol 2:395–411.

49. Nair R, et al. (2014) Extracellular vesicles derived from preosteoblasts influence embryonic stem cell differentiation. Stem Cells Dev 23:1625–1635.

50. Lawson J, Dickman C, MacLellan S, Towle R, Jabalee J, Lam S & Garnis C (2017) Selective secretion of microRNAs from lung cancer cells via extracellular vesicles promotes CAMK1D-mediated tube formation in endothelial cells. Oncotarget 8:83913–83924.

51. Veziroglu EM & Mias GI (2020) Characterizing Extracellular Vesicles and Their Diverse RNA Contents. Front Genet 11:700.

52. Srinivasan S et al. (2019) Small RNA Sequencing across diverse biofluids identifies optimal methods for exRNA isolation. Cell 177:446–462.

53. Tosar JP, Cayota A (2020) Extracellular tRNAs and tRNA-derived fragments. RNA Biol 17:1149–1167.

54. Vidal M (2019) Exosomes: Revisiting their role as “garbage bags”. Traffic 20:815–828.

55. Hulstaert E et al. (2020) Charting extracellular transcriptomes in the human biofluid RNA atlas. Cell Rep 33:108552.

56. Martin M (2011) Cutadapt removes adapter sequences from high-throughput sequencing reads. EMBnet J 17:10–12. DOI: https://doi.org/10.14806/ej.17.1.200

57. Langmead B, Trapnell C, Pop M, & Salzberg SL (2009) Ultrafast and memory-efficient alignment of short DNA sequences to the human genome. Genome Biol 10:R25.

58. Love MI, Huber W, & Anders S (2014) Moderated estimation of fold change and dispersion for RNA-seq data with DESeq2. Genome Biol 15:550.

59. Holmes AD, Howard JM, Chan PP, & Lowe TM (2020) tRNA Analysis of eXpression (tRAX): A tool for integrating analysis of tRNAs, tRNA-derived small RNAs, and tRNA modifications. Submitted. http://trna.ucsc.edu/tRAX/citation/

60. Friedländer MR, Mackowiak SD, Li N, Chen W, & Rajewsky N (2012) miRDeep2 accurately identifies known and hundreds of novel microRNA genes in seven animal clades. Nucl Acids Res 40:37–52.

61. Li B & Dewey CN (2011) RSEM: accurate transcript quantification from RNA-Seq data with or without a reference genome. BMC Bioinformatics 12:323.

